# HIF-1–regulated TPM3 links hypoxia to motility and invasion beyond the hypoxic fraction in triple-negative breast cancer

**DOI:** 10.64898/2026.01.08.698356

**Authors:** Chumin Zhou, Jack T. Crusher, Kate Friesen, Sophie A. Twigger, Eduard Petrosyan, Graham Booker, Priya Samuel, Eileen E. Parkes, Ester M. Hammond

**Affiliations:** Department of Oncology, University of Oxford, Oxford, OX3 7DQ, UK; School of Biological & Medical Sciences, Oxford Brookes University, Headington, Oxford OX3 0BP, UK

**Keywords:** Hypoxia, TNBC, TPM3, motility, Extracellular Vesicles

## Abstract

Hypoxia is a defining feature of triple-negative breast cancer (TNBC), driving invasion, metastasis, and therapy resistance. Understanding the molecular effectors of hypoxia is essential to identify new therapeutic targets. Here, we investigated tropomyosin 3 (TPM3), an actin-binding protein that regulates filament stability. TPM3 is significantly upregulated in breast cancer, including in TNBC, where elevated levels correlate with poor overall survival. Using validated hypoxia signatures and TNBC cell models, we show that TPM3 is induced in physiologically relevant hypoxic conditions in a HIF-1–dependent manner. Both mRNA and protein levels of TPM3 increased in response to hypoxia, and TPM3 colocalised with F-actin, supporting cytoskeletal organisation. Functional assays demonstrated that depletion or inhibition of TPM3 impaired cell morphology, motility, and invasion in hypoxic TNBC cells, while not affecting viability. Notably, TPM3 inhibition synergised with Paclitaxel and Doxorubicin, enhancing therapeutic efficacy. In addition, TPM3 was incorporated into extracellular vesicles (EVs), with hypoxia increasing EV-mediated transfer of TPM3 to normoxic cells and promoting their motility. These findings establish TPM3 as a hypoxia-inducible, HIF-1–regulated effector of cytoskeletal dynamics and intercellular communication, underscoring its potential as a therapeutic target to limit TNBC aggressiveness and improve treatment outcomes.

## Introduction

Hypoxia (conditions of insufficient oxygen) is a hallmark of the tumour microenvironment (TME) in solid cancers, arising as a consequence of rapid tumour growth and inadequate or inefficient vasculature. Hypoxia is associated with therapy resistance and poor patient prognosis ^1^. The adverse prognostic impact of hypoxia has been particularly well documented in breast cancer and is even more pronounced in triple-negative breast cancer (TNBC), the most aggressive subtype which lacks targeted treatment options ^2,3^. Patients who succumb to TNBC do so because of metastatic disease and while a relationship between hypoxia and increased metastasis is well-known, the underpinning mechanisms are less clear. Emerging evidence suggests that hypoxia-driven changes in gene expression, activation of hypoxia-inducible factors (HIFs), epithelial-to-mesenchymal transition (EMT), metabolic reprogramming, and remodelling of the TME may all contribute to enhanced metastatic dissemination ^4,5^. The identification of hypoxic tumour markers which predict poor patient outcome is therefore critical to identify novel therapeutic strategies.

TPM3 (Tropomyosin 3) encodes a member of the tropomyosin family of actin-binding proteins which stabilise actin filaments and regulate the cytoskeleton. TPM3 plays important roles in maintaining cell structure, intracellular transport, and muscle contraction ^6,7^. While TPM3 function has been predominantly characterised in muscle cells/myopathies, dysregulation or gene fusions involving TPM3 have been implicated in various cancers ^7–13^. The mRNA for TPM3 has been described in microvesicles generated by platelets and it has been suggested that TPM3 could be delivered to breast cancer cells and promote metastasis ^14^. Interestingly, the hypoxia-inducible Right Open Reading Frame Kinase 3 (RIOK3) was found to interact with TPM3 leading to the hypothesis that this interaction could support RIOK3-dependent migration and invasion in hypoxia ^15^. Together, these studies led us to investigate a role for TPM3 in the biological response to hypoxia.

Here, we found that TPM3 is a target of the HIF-1 transcription factor in a broad range of hypoxic conditions and controls the motility and invasion capacity of hypoxic TNBC cells. Most importantly, we identify TPM3 as an extracellular vesicle (EV) cargo protein which can increase the motility of normoxic cells. Together, these data suggest that the hypoxia-mediated induction of TPM3 contributes to the metastatic potential of both the hypoxic and oxygenated tumour fractions of TNBC.

## Results

### TPM3 is hypoxia-inducible in conditions relevant to TNBC

TPM3 expression was found to be significantly higher in breast cancer (BRCA) tissue, compared to normal breast tissue (**Figure 1A** and **Figure S1A**). Furthermore, TPM3 expression was found to be significantly higher in TNBC compared to normal tissue (**Figure 1B** and **Figure S1B**). Next, we asked how TPM3 expression correlated with TNBC patient survival and found that high TPM3 is associated with a poorer overall survival rate in TNBC (**Figure 1C**). To further explore a relationship between TPM3 and hypoxia, we used two validated hypoxia signatures and found a significant correlation in patient samples, demonstrating that TPM3 expression is increased in more hypoxic TNBC (**Figure 1D** and S1C) 16,17.

**Figure 1.**
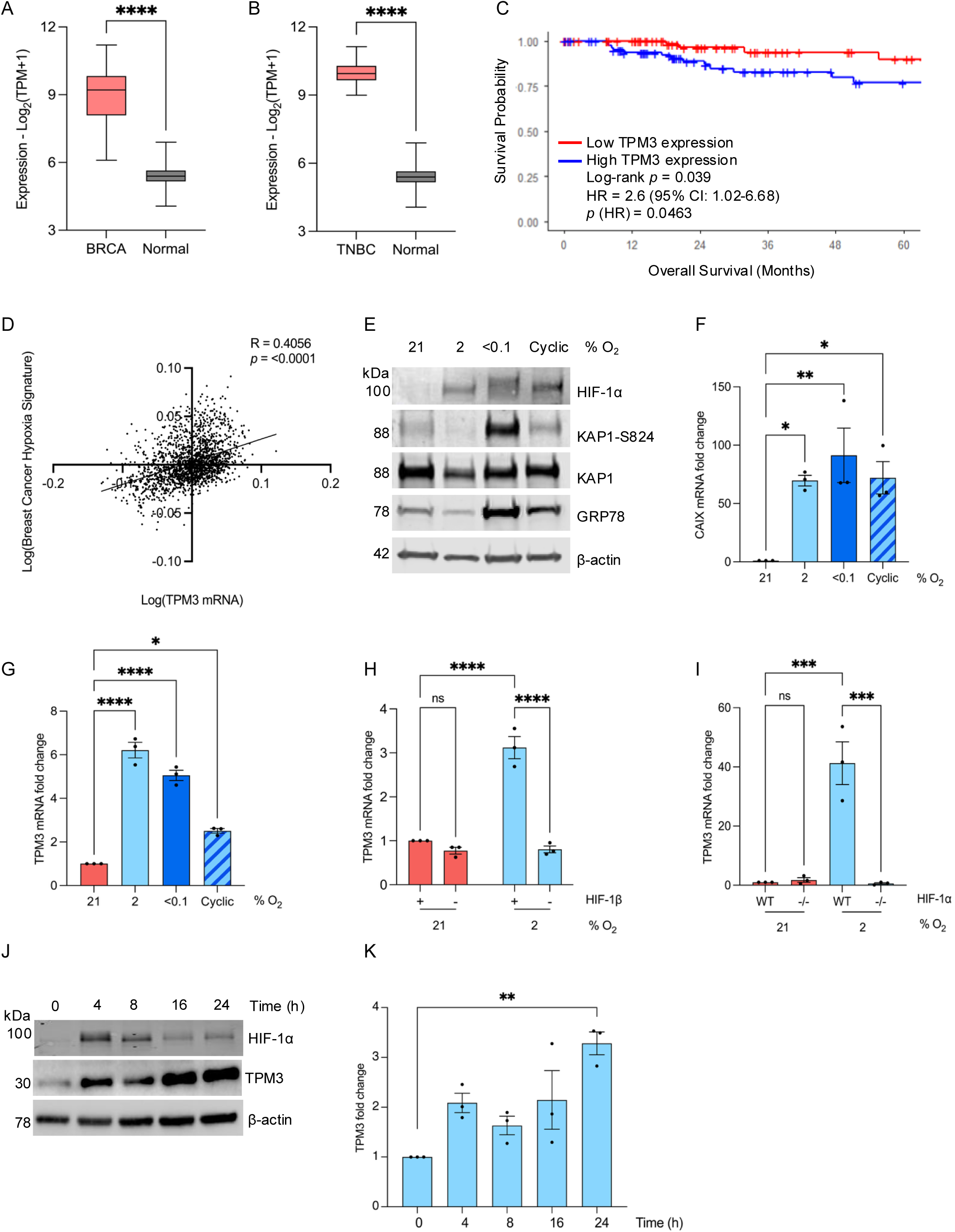
TPM3 is induced in hypoxia in a HIF-1-dependent manner. **A.** TPM3 mRNA levels in BRCA and normal breast tissue generated using TPM3 mRNA expression from TCGA-BRCA. num(BRCA)=1082; num(Normal)=514 **B.** TPM3 mRNA levels in TNBC and normal breast tissue generated using TPM3 mRNA from Genotype-Tissue Expression (GTEx). num(TNBC)=171; num(Normal)=514 **C.** Kaplan-Meier curve of overall survival in TNBC patients with high or low TPM3 expression, generated using TCGA-BRCA (dichotomised at the median). Statistical significance was assessed by log-rank test. **D.** Correlation between the Buffa hypoxia signature and TPM3 mRNA expression from Metabric dataset. Statistical analysis was determined using simple linear regression test and Pearson correlation. **E.** MDA-MB-231 cells were exposed to a range of hypoxic conditions for 16 h followed by western blotting for the proteins indicated, β-actin was used as a loading control. **F.** MDA-MB-231 cells were exposed to a range of hypoxic conditions for 16 h followed by RT-qPCR to determine CAIX mRNA level. Statistical testing was done using a paired *t*-test. **G.** MDA-MB-231 cells were exposed to a range of hypoxic conditions for 16 h followed by RT-qPCR for TPM3. Statistical testing was done using a paired *t*-test. **H.** MDA-MB-231 cells were treated with siRNA to HIF-1β followed by exposure to 21% or 2% O_2_ for 16 h. TPM3 mRNA level was determined and shown relative to the normoxic control. Statistical testing was done by two-way ANOVA with Šídák’s multiple comparisons test. **I.** RKO and RKO^HIF-11l!-/-^ cells were exposed to hypoxia (2% O_2_) for 16 h followed by RT-qPCR for TPM3. Statistical testing was done by two-way ANOVA with Tukey’s multiple comparisons test. **J.** MDA-MB-231 cells were exposed to 2% O_2_ for the times shown followed by western blotting for the proteins indicated. **K.** Quantification of TPM3 levels in cells treated as in part J. Statistical testing was done by paired *t*-test. Data shown from three separate experiments (*n* = 3) are displayed with mean ± standard error of the mean (SEM) unless specified otherwise. Statistical testing was carried out as indicated. * *p* < 0.05, ** *p* < 0.01, *** *p* < 0.001, **** *p* < 0.0001 and ns *p* > 0.05.

Tumour hypoxia exists as a gradient of oxygen tensions within a tumour and also includes transitions between levels, known as cyclic or intermittent hypoxia ^18^. It is important to consider a range of hypoxic conditions when investigating hypoxia-mediated biology as the biological response can differ ^19^. Here, we considered 2% O_2_, <0.1% O_2_ and cyclic conditions (transitions between 2 and <0.1% O_2_). As expected, HIF-1α was stabilised in all of the hypoxic conditions, however evidence of the DNA damage response was limited to <0.1% O_2_ and the unfolded protein response to <0.1% O_2_ and cyclic conditions (**Figure 1E**)^20^. TNBC cell lines (MDA-MB-231 and MDA-MB-453) were exposed to the hypoxic conditions (2, <0.1% O_2_ and cyclic) and changes in TPM3 mRNA determined. As expected, the well-validated HIF target, CAIX was induced in response to hypoxia (**Figure 1F**) ^21^. A significant increase in TPM3 mRNA was observed in all the hypoxic conditions tested in both cell lines demonstrating that TPM3 is hypoxia inducible in a broad range of hypoxic conditions relevant to TNBC (**Figure 1G** and **S1D**).

This hypoxia-mediated induction in all three conditions and correlation with hypoxia signatures suggested the possibility that TPM3 was regulated by one of the hypoxia inducible factors (HIFs). In support of this, a hypoxia responsive element (HRE) was identified upstream of the TPM3 coding sequence (**Figure S1E**). To investigate possible HIF dependence, we treated MDA-MB-231 cells with siRNA to HIF-1β to prevent both HIF-1 and HIF-2 mediated signalling and determined the impact on TPM3 expression. Loss of HIF-1β abrogated the induction of TPM3 and CAIX in hypoxia (**Figure 1H** and **S1F-H**). To determine if TPM3 induction in hypoxia was mediated by HIF-1 or HIF-2, we used the matched colorectal cell lines RKO and RKO^HIF-1^^α-/-^ and found that TPM3 induction in hypoxia was abrogated by loss of HIF-1α (**Figure 1I** and **S1I-K**). Notably, these data do not rule out the possibility that HIF-2 could also regulate TPM3 expression at specific oxygen levels or times. Together, these data confirm that TPM3 is hypoxia-inducible in all the cell lines investigated and suggests that this occurs in a HIF-1-dependent manner.

MDA-MB-231 and MDA-MB-453 cells were then exposed to hypoxia (2% O_2_) and western blotting carried out for TPM3. TPM3 protein was found to increase in response to hypoxia in both cell lines (MDA-MB-231 **Figure 1J, K** and MDA-MB-453 **S2A, B**). Hypoxia-mediated induction of TPM3 was further confirmed in response to <0.1% O_2_, cyclic conditions and a third TNBC cell line (BT-549) (**Figure S2C-E**).

### TPM3 promotes motility and invasion in hypoxia

Next, we validated that TPM3 colocalised with F-actin (detected using phalloidin) ^22^. TPM3 appeared predominantly cytoplasmic with an apparent filamentous structure. Notably, a punctate signal near to the nucleus was also observed however this was unaffected by siRNA mediated knockdown of TPM3 and was therefore considered non-specific staining (**Figure S2F-G**). A clear colocalisation of TPM3 and F-actin was observed which did not change in response to hypoxia (**Figure 2A, B**). Next, we analysed cell morphology with and without siTPM3 knockdown and observed an increase in elongated trailing edges following loss of TPM3, suggesting a loss of actin filament structure and a reduced ability to contract the trailing edge during migration. To confirm, we measured the circularity of cells and found a significant reduction in cell circularity when TPM3 was depleted, in both normoxic and hypoxic (2% O_2_) conditions (**Figure 2C**). As loss of TPM3 appeared to impair the ability of MDA-MB-231 cells to recoil their trailing edge, we hypothesised that TPM3 may also influence the leading edge, where F-actin stabilisation in lamellipodia and focal adhesions is essential for generating the mechanical force required for trailing edge recoil ^23^. Loss of TPM3 significantly reduced the intensity of phalloidin staining at the leading edge under both hypoxic (2% O_2_) and normoxic conditions (**Figure 2D**). Depletion of TPM3 had no effect on the width of the leading edge under normoxia but had a significant effect under hypoxia, further supporting a role for TPM3 in stabilising F-actin and cell motility (**Figure 2E, F**). Together, our data suggest the hypothesis that TPM3, through its role in stabilising actin filaments, could promote the motility of hypoxic cells and therefore drive metastatic progression in aggressive TNBC.

**Figure 2.**
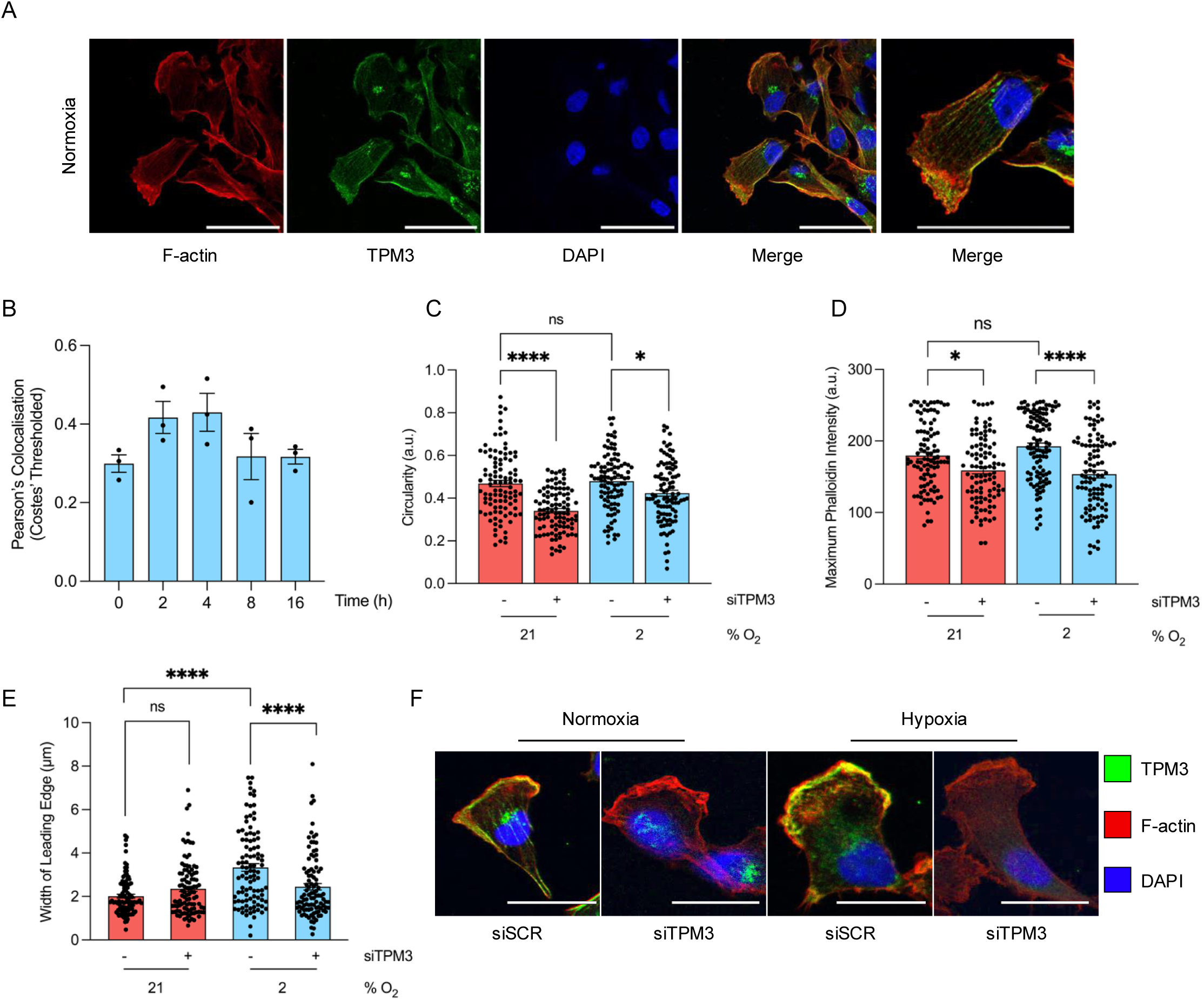
TPM3 affects F-actin organisation at the leading edge under hypoxia. **A.** Representative images of immunofluorescence of MDA-MB-231 cells. Cells were stained for F-actin (phalloidin; red), TPM3 (green) and DAPI (blue). Scale bar is 50 μm. **B.** MDA-MB-231 cells were exposed to hypoxia (2% O_2_) for the times indicated followed by staining for TPM3 and F-actin. Quantification of colocalisation of TPM3 and F-actin was carried out using Costes’ thresholded Pearson’s coefficient. **C.** MDA-MB-231 cells were treated with a scramble siRNA or siTPM3 and then exposed to 21% or 2% O_2_ for 16 h. Cells were then stained for TPM3, F-actin and DAPI. Quantification of cell circularity (a.u.) was determined and shown. Data collected from ≥100 cells per condition. Statistical testing was done by one-way ANOVA with Tukey’s multiple comparisons test. **D.** Cells were treated as in part A. Quantification of maximum phalloidin intensity at the leading edge was determined. Data collected from ≥100 cells per condition. Statistical testing was done by one-way ANOVA with Tukey’s multiple comparisons test. **E.** Cells treated as in part A. Quantification of the leading-edge width is shown. Data collected from ≥100 cells per condition. Statistical testing was done by one-way ANOVA with Tukey’s multiple comparisons test. **F.** Representative immunofluorescence microscopy images showing TPM3 (green), F-actin (phalloidin; red), and DAPI (blue) at the leading edge of MDA-MB-231 cells, transfected with scramble control (siSCR) or TPM3 siRNA in 21% or 2% O_2_. Scale bar is 50 μm. Data shown from three separate experiments (*n* = 3) are displayed with mean ± standard error of the mean (SEM) unless specified otherwise. * *p* < 0.05, **** *p* < 0.0001 and ns *p* > 0.05.

To test the role of TPM3 in hypoxia-mediated motility, we carried out wound healing assays in MDA-MB-231 cells with siRNA-mediated depletion of TPM3. Before investigating motility, we verified that cell cycle distribution and viability were not altered in the hypoxic conditions used (**Figure S3A-C**). Furthermore, we demonstrated that loss of TPM3 had no significant effect on clonogenic survival of cells in any of the hypoxic conditions tested (**Figure S3D-E**). Depletion of TPM3 was found to significantly slow down wound closure in hypoxic conditions but had no impact in normoxia (**Figure 3A-C** and **S4A, B**). A small molecule inhibitor of TPM3 has been described (ATM-3507) which we also used to test the role of TPM3 in hypoxia-mediated motility ^24^. Again, we first verified that ATM-3507 did not significantly impact viability in both normoxic and hypoxic conditions at the dose reported to inhibit TPM3 activity (**Figure S4C**) ^12^. Cells exposed to ATM-3507 were significantly less motile in hypoxic conditions (**Figure 3D, E** and **S4D, E**). Both motility and the ability to invade play important roles in metastatic potential, therefore we investigated the contribution of TPM3 to hypoxia-mediated invasion ^25^. As expected, hypoxic cells showed a greater invasion capacity compared to the normoxic however, this was significantly reduced when TPM3 was depleted (**Figure 3F-H** and **S4F, G**). In conclusion, TPM3 has an important role in cell migration and invasion in hypoxic TNBC cells.

**Figure 3.**
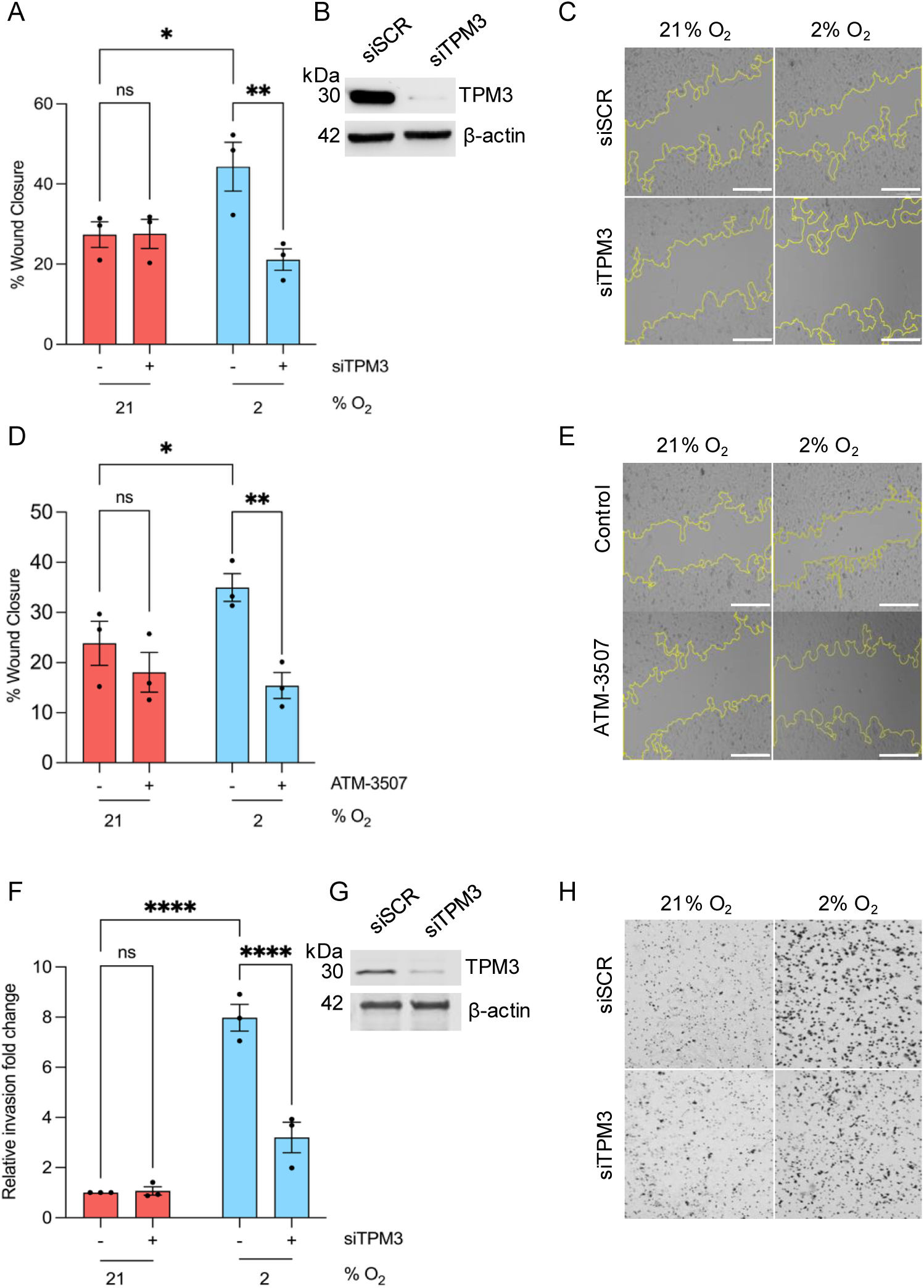
Hypoxia-mediated migration and invasion are TPM3 dependent. **A.** MDA-MB-231 cells were treated with siTPM3 or a scramble siRNA followed by exposure to 21% or 2% O_2_ for 8 h. The % of wound closure is shown in each condition. Statistical testing was done by two-way ANOVA with Šídák’s multiple comparison test. **B.** Knockdown of TPM3 in cells part A was validated by western blotting, β-actin was used as a loading control. **C.** Representative images of wound healing assay in part A. Scale bars are 200 μm. **D.** MDA-MB-231 cells were treated with ATM-3507 (6 µM) for 1 h prior to and during exposure to 21% or 2% O_2_ for 8 h. The % of wound closure is shown in each condition. Statistical testing was done using a two-way ANOVA with Šídák’s multiple comparison test. **E.** Representative images of wound healing assay in part D. Scale bars are 200 μm. **F.** MDA-MB-231 cells were treated with siTPM3 or a scramble siRNA (siSCR) followed by exposure to 21% or 2% O_2_ for 16 h. The invasion fold change relative to normoxic control is shown in each condition. Statistical testing was done by two-way ANOVA with Šídák’s multiple comparison test. **G.** Knockdown of TPM3 in cells used in part F was validated by western blotting, β-actin was used as a loading control. **H.** Representative images of invasion assay in part F. Data from three separate experiments (*n* = 3) are displayed with mean ± standard error of the mean (SEM) unless specified otherwise. * *p* < 0.05, ** *p* < 0.01, **** *p* < 0.0001 and ns *p* > 0.05.

### Combination of TPM3 inhibition with TNBC standard of care

The standard of care for TNBC patients includes Carboplatin, Doxorubicin, Paclitaxel and radiotherapy. Inhibition of TPM3 has been shown to combine effectively with Paclitaxel and Doxorubicin in normoxia, resulting in the reduction of cell viability in neuroblastoma and ovarian cancer cell lines ^12,26^. Here, we investigated the efficacy of combining TPM3 inhibition or siRNA-mediated loss with the standard of care for TNBC in physiologically relevant conditions. MDA-MB-231 cells were exposed to hypoxia (2% O_2_) and a range of doses of ATM-3507 with Carboplatin, Doxorubicin or Paclitaxel followed by an assay for viability. The Highest Single Agent (HSA) model was used to investigate potential synergy and revealed a reduction of cell viability across increasing concentrations of ATM-3507, Carboplatin, Doxorubicin and Paclitaxel (**Figure S5A-C**). In addition, strong synergy (score >10) was observed with ATM-3507 combined with Doxorubicin or Paclitaxel, no antagonistic or additivity regions were indicated (**Figure 4A, B, C**). To determine any effect on radiosensitivity, MDA-MB-231 cells were treated with siTPM3 followed by irradiation in hypoxia (2% O_2_). Loss of TPM3 had no impact on radiosensitivity in normoxia or hypoxia (**Figure 4D** and **S5D**). In addition, inhibition of TPM3 in normoxia or hypoxia (2% O_2_) did not significantly impact radiosensitivity (**Figure S5E**). Together, these data suggest that inhibition of TPM3 could be combined with standard of care for TNBC to potentially reduce metastatic spread and improve patient prognosis.

**Figure 4.**
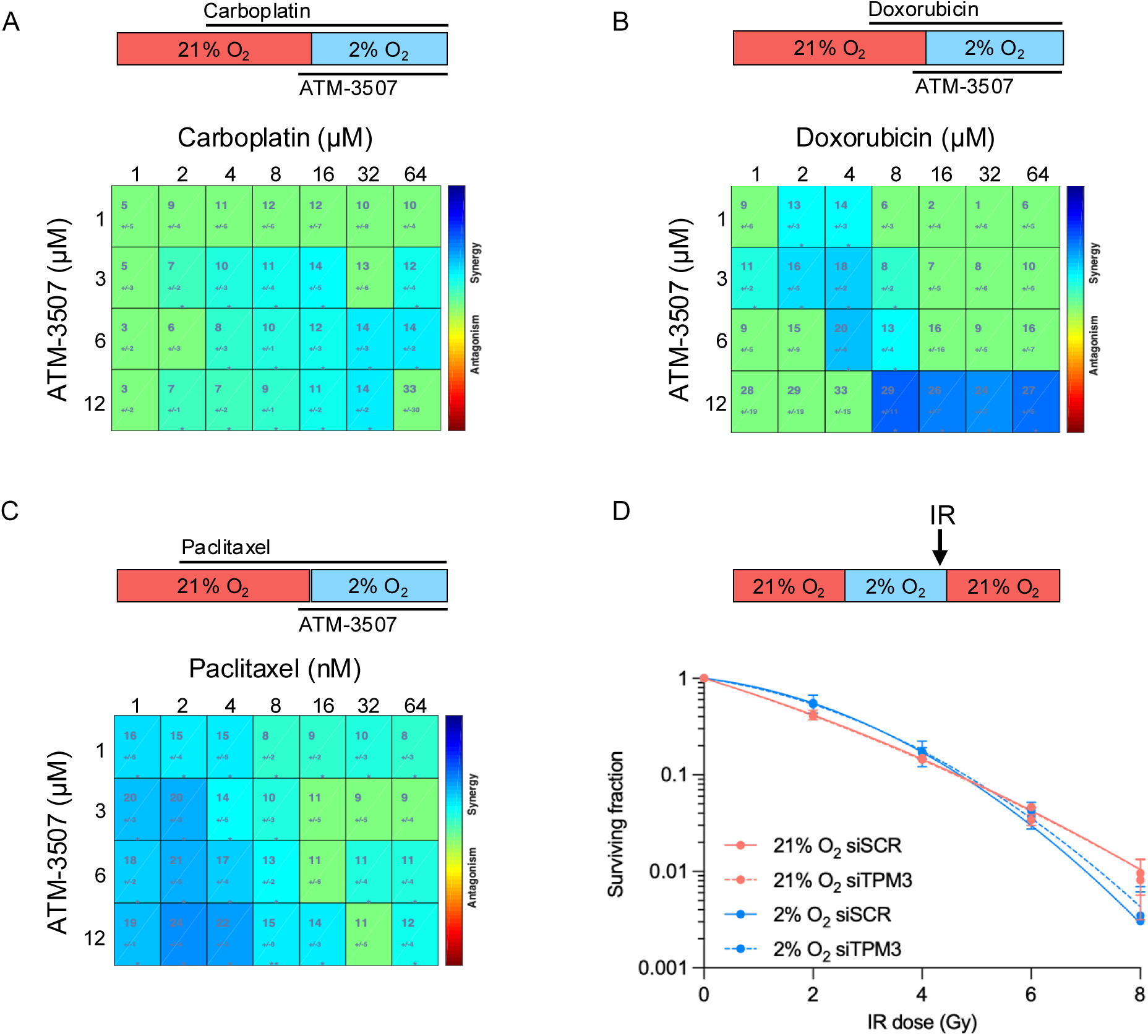
Inhibition of TPM3 combines effectively with standard of care for TNBC. **A.** MDA-MB-231 cells were treated with Carboplatin (1 - 64 μM) for 48 h and ATM-3507 (1.5-12 μM) for 17 h. For the 16 h before an MTT assay was carried out the cells were in hypoxia (2% O_2_). **B.** MDA-MB-231 cells were treated with Doxorubicin (1 - 64 μM) for 24 h and ATM-3507 (1.5-12 μM) for 17 h. For the 16 h before an MTT assay was carried out the cells were in hypoxia (2% O_2_). **C.** MDA-MB-231 cells were treated with Paclitaxel (1 - 64 nM) for 72 h and ATM-3507 (1.5-12 μM) for 17 h. For the 16 h before an MTT assay was carried out the cells were in hypoxia (2% O_2_). In A, B and C Highest Single Agent (HSA) synergy score was assessed by an interactive platform Combenefit, scores > 0 represent synergism (blue) and scores < 0 represent antagonism (red). Dose-response matrices with drug concentrations on axes and combination effects as a heatmap overlay are shown. Statistical testing was carried out using the built-in analysis algorithm ^39^. **D.** MDA-MB-231 cells were treated with siTPM3 or a scramble siRNA (siSCR) followed by exposure to 21% or 2% O_2_ for 16 h prior to irradiation using the doses indicated. Cells in hypoxia were irradiated in hypoxic conditions (shown schematically). After irradiation, all cells were returned to 21% O_2_ and a colony survival assay carried out. Data from three separate experiments (*n* = 3) are displayed with mean ± standard error of the mean (SEM).

### Hypoxia-induced TPM3 influences motility in normoxic cells

The hypoxic TME is known to impact neighbouring oxygenated cancer cells although a role for TPM3 in this has not been described ^27^. MDA-MB-231 cells were treated with siRNA TPM3, exposed to normoxia or hypoxia (2% O_2_) and conditioned media collected and transferred to normoxic cells followed by a wound healing assay (**Figure 5A, B**). Conditioned media from hypoxic cells (donor cells) when added to normoxic cells (recipient cells) significantly increased the wound healing rate of the normoxic cells. However, when the conditioned media from TPM3 depleted donor cells was added to recipient cells no increase in the motility of the normoxic recipients was observed (**Figure 5C, D**). This finding suggests that TPM3 contributes to factors present in the conditioned media or regulates the export of proteins and/or EVs from hypoxic cells, thereby influencing the migratory behaviour of normoxic recipient cells.

**Figure 5.**
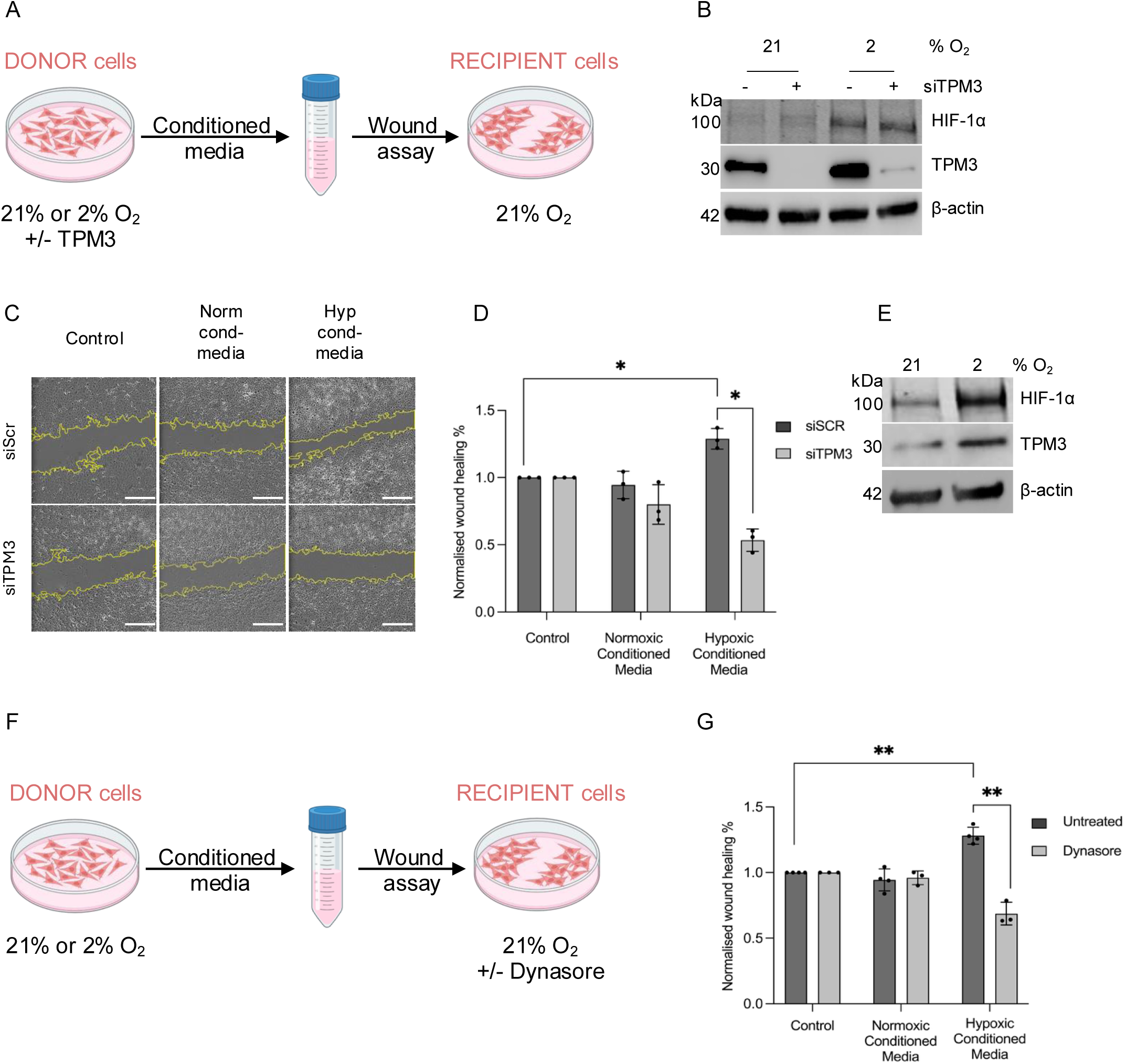
Conditioned media from hypoxic cells increase migration in normoxic cells in a TPM3-dependent manner. **A.** Schematic representation of the wound assay used to determine the impact of TPM3 loss in donor cells on recipient cell migration. **B.** MDA-MB-231 (donor) cells were treated with siTPM3 or a scramble siRNA and exposed to 21% or 2% O_2_ for 16 h, followed by western blotting to validate TPM3 knockdown and HIF-1α stabilisation. **C.** Representative images of wound healing assay from cells treated as in part D. Scale bars, 500 μm. **D.** MDA-MB-231 donor cells were treated with siTPM3 or a scramble siRNA (siSCR) followed by exposure to 21% or 2% O_2_ for 16 h. Conditioned media was then collected and applied to wounded MDA-MB-231 recipient cells in 21% O_2_ for 16 h. Quantification of normalised wound healing % is shown as fold change relative to control media. Statistical testing was done by paired *t*-test. **E.** MDA-MB-231 donor cells were exposed to 21% and 2% O_2_ for 16 h, followed by western blotting to indicated proteins. **F.** Schematic representation of the wound healing assay used to determine the impact of Dynasore. **G.** MDA-MB-231 donor cells were exposed to 21% or 2% O_2_ for 16 h. Conditioned media was then collected and added to wounded MDA-MB-231 normoxic recipient cells +/-Dynasore (50 μM) for 16 h. Quantification of normalised wound healing % is shown as fold change relative to control media. Statistical testing was done by paired *t*-test. Data from three separate experiments (*n* = 3) are displayed with mean ± standard error of the mean (SEM). * *p* < 0.05, ** *p* < 0.01.

To investigate further, we again generated conditioned media from normoxic and hypoxic (2% O_2_) cells and added it to wounded normoxic recipient cells (**Figure 5E**). However, this time we also included the dynamin inhibitor, Dynasore, to prevent EV uptake in the recipient cells (**Figure 5F**). As previously, we saw a significant increase in wound healing with the addition of hypoxic conditioned media. However, when Dynasore was added a significant reduction in wound closure rate was observed (**Figure 5F, G** and **S6A**). This suggests EV uptake from the hypoxic conditioned media is critical to the impact on recipient cell motility in normoxia. To account for the known off-target effect of Dynasore in reducing basal motility, data were normalised to control cells treated with Dynasore alone ^28^. Importantly, we confirmed that Dynasore did not affect TPM3 expression validating that the reduction of wound healing in hypoxic conditioned media following Dynasore treatment was not due to TPM3 suppression in the migrating cells (**Figure S6B**). These data again confirm that hypoxia-induced TPM3 contributes to a conditioned media which has the capacity to increase the motility of normoxic cells and implicates EV transfer as the underlying mechanism. Therefore, we isolated and purified EVs to confirm if they were responsible for the impact on normoxic cell motility.

First, we verified using transmission electron microscopy (TEM) that EV production could be seen in MDA-MB-231 cells in hypoxic conditions. Late endosomes and EVs were observed in normoxia and hypoxia with an apparent increase in vesicle release in hypoxia (2% O_2_) (**Figure 6A**). A clear increase in the abundance of mitochondria following 2% O_2_ exposure was also observed and has been reported previously ^29^. To investigate the apparent increase in EVs detected by TEM, we isolated EVs from the conditioned media of normoxic and hypoxic (2% O_2_) MDA-MB-231 cells and quantified them using nanoparticle tracking analysis (NTA). The total concentration of EVs increased significantly in hypoxia (2% O_2_), while the size distribution was similar in normoxic and hypoxic conditions (**Figure 6B-D**). Next, to determine if EVs released by hypoxic cells impacted recipient cell migration, MDA-MB-231 cells were again exposed to hypoxia (2% O_2_) and TPM3 induction confirmed (**Figure 6E**). EVs were then isolated and added to wounded normoxic recipient cells to determine impact on wound healing rate (**Figure 6F**). EVs isolated from hypoxic cells increased the motility of normoxic cells and this was abrogated in the presence of Dynasore (**Figure 6G** and **S6C**).

**Figure 6.**
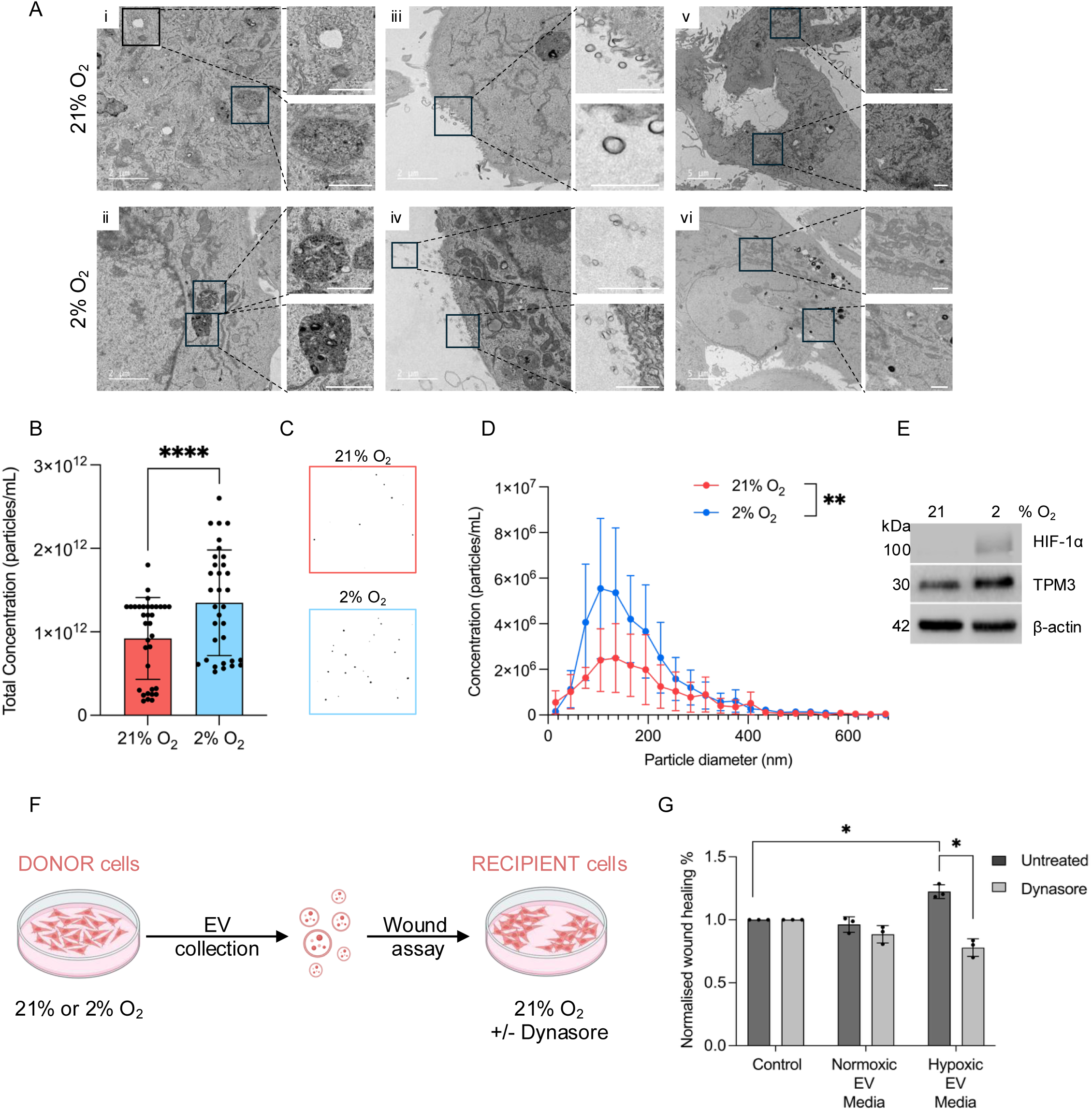
EVs generated by hypoxic cells increase migration of normoxic cells. **A.** MDA-MB-231 cells in 21% O_2_ and 2% O_2_ (12 h) were processed and imaged by TEM. Images of late endosomes (enlarged in i, ii), EVs (enlarged in iii, iv) and mitochondria (enlarged in v, vi) are shown. Scale bars are indicated on each image. **B.** MDA-MB-231 cells were exposed to 21% or 2% O_2_ for 24 h followed by EV isolation. Nanoparticle tracking analysis (NTA) was then carried out to determine the total concentration of EVs. Statistical testing was done by paired *t*-test. **C.** Representative images of NTA from part B. **D.** Particle diameter was determined for the EVs isolated in part B using NTA. **E.** MDA-MB-231 cells were exposed to 21% or 2% O_2_ for 24 h, followed by western blotting of whole cell lysates for the indicated proteins. **F.** Schematic representation of the wound healing assay to determine the impact of addition of EVs from normoxic or hypoxic cells on normoxic recipient cells. **G.** MDA-MB-231 donor cells were exposed to 21% or 2% O_2_ for 24 h. Conditioned media was then collected and EVs isolated. EVs were then resuspended in FBS free normoxic culture media and added to MDA-MB-231 normoxic recipient cells +/- Dynasore (50 μM) for 16 h. Quantification of normalised wound healing % is shown as fold change relative to control media. Statistical testing was done by paired *t*-test. Data from three separate experiments (*n* = 3) are displayed with mean ± standard error of the mean (SEM). * *p* < 0.05, ** *p* < 0.01, and **** *p* < 0.0001.

### TPM3 is an EV cargo protein

These data led us to the non-mutually exclusive conclusions that TPM3 could play a role in EV production in hypoxic cells or that TPM3 could be transferred as cargo through EVs to recipient cells. MDA-MB-231 cells were treated with scramble siRNA or siTPM3 and exposed to hypoxia (2% O_2_) followed by EV isolation and NTA. Again, EV production was increased in hypoxic conditions but loss of TPM3 did not significantly change EV production or size distribution in normoxia or hypoxia (**Figure 7A, B, C**). Next, we again isolated EVs from normoxic and hypoxic cells and used NTA to ensure equal loading for western blotting. ALIX is a well-characterised EV protein and was used as a control ^30^. Blotting for the mitochondrial protein PRDX3 and Golgi-associated GM130 was also included to demonstrate the purity of the EVs ^31^. As expected, western blotting of whole cell lysates showed that ALIX, PRDX3 and GM130 were expressed equally in normoxic and hypoxic cells while TPM3 was induced in hypoxia (**Figure 7D**). TPM3 was found to be included in normoxic EVs and the levels increased in EVs generated from hypoxic cells therefore validating TPM3 as an EV cargo protein (**Figure 7E, F**).

**Figure 7.**
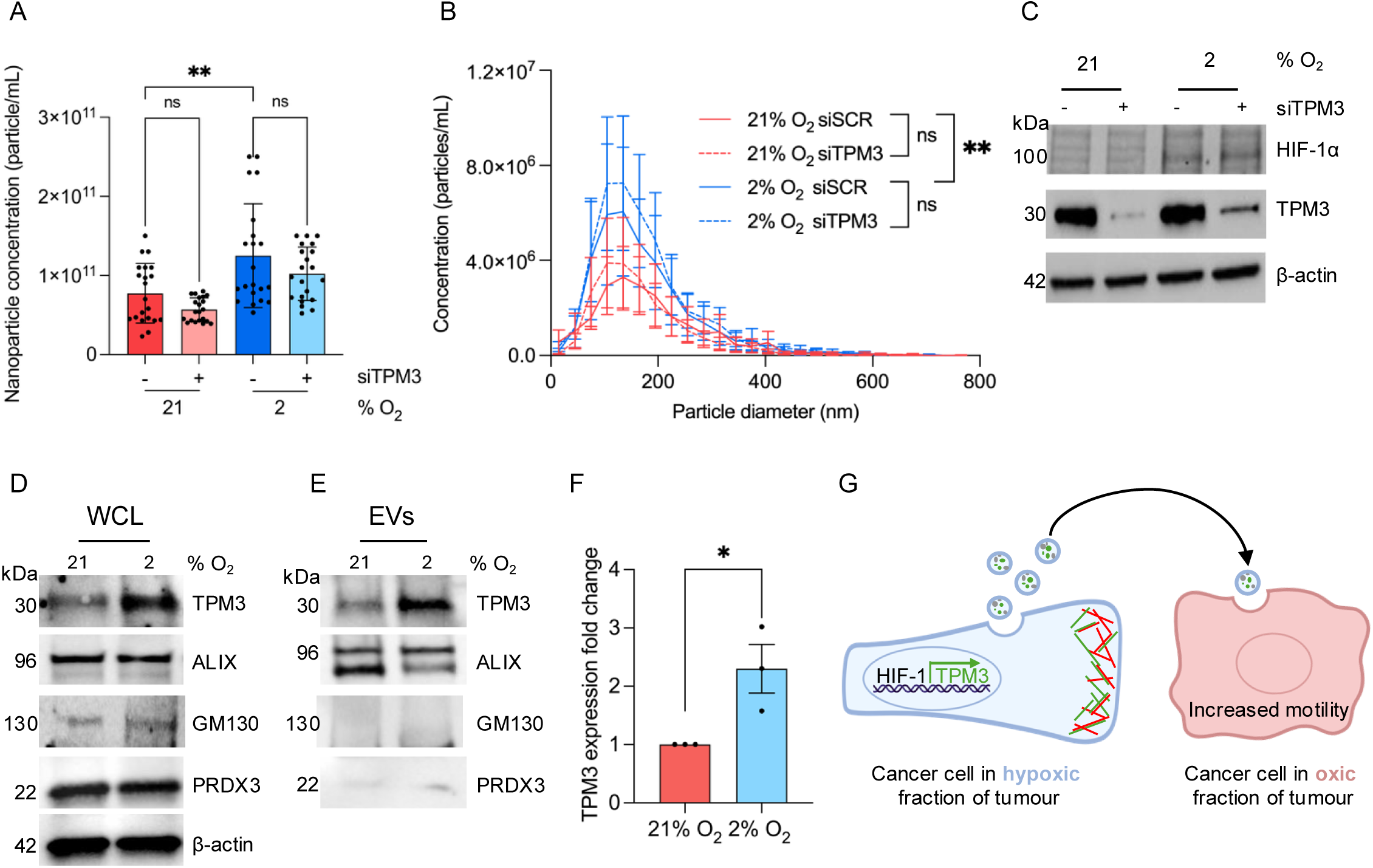
TPM3 is an EV cargo protein. **A.** MDA-MB-231 cells were treated with siTPM3 or a scramble siRNA and exposed to 21% or 2% O_2_ for 24 h. EVs were isolated, followed by nanoparticle tracking analysis (NTA). The total concentration of EVs from each condition is shown. Statistical testing was done by paired *t*-test. **B.** Particle diameter distribution of the EVs isolated in part A. Statistical testing was done using a paired *t*-test. **C.** Western blotting of lysates from cells in part A to verify TPM3 knockdown and HIF-1α stabilisation. **D.** MDA-MB-231 cells cultured under normoxic (21% O_2_) or hypoxic (2% O_2_) conditions for 24 h followed by western blotting of whole cell lysates (WCL) as indicated. **E.** Western blotting of purified EVs isolated from cells in part D followed by NTA quantification to ensure equal loading. ALIX was used as an EV marker and PRDX3/GM130 used to validate EV purity. **F.** Quantification of TPM3 protein expression in EVs from part E. TPM3 band intensity was normalised to ALIX (loading control) and expressed relative to 21% O_2_. **G.** A summary schematic. TPM3 is induced in a broad range of physiologically relevant hypoxic conditions in a HIF-1 dependent manner. The accumulation of TPM3 (shown in green) contributes to F-actin (shown in red) stabilisation at the leading edge and increased cell motility/invasion. EVs produced by the hypoxic cells contain TPM3, in addition to many other proteins, and when transferred to cells in normoxia (oxic) conditions also increases their motility. * *p* < 0.05, ** *p* < 0.01, and ns *p* > 0.05.

## Discussion

In this study, we demonstrate that TPM3 is a hypoxia-inducible gene in TNBC, and that this occurs in a HIF-1-dependent manner. Under hypoxic conditions, TPM3 supports actin filament stability, maintaining cell shape and enabling efficient migration and invasion. Loss of TPM3 disrupted F-actin organisation at both the leading and trailing edges, reduced cell circularity, and impaired motility. Importantly, TPM3 is also incorporated into EVs released by hypoxic cells, which enhance motility in normoxic recipient cells. Together, these findings demonstrate that hypoxia-induced TPM3 enhances migration capacity across the tumour, extending its impact beyond the hypoxic fraction (**Figure 7G**). Our findings support that targeting TPM3 during treatment of TNBC could reduce metastatic burden, by reducing the migratory potential and invasiveness of residual hypoxic and normoxic cancer cells. Importantly, TPM3 is druggable and appeared well-tolerated *in vivo* as evidenced by preclinical studies with ATM-3507 and its precursor TR100 ^12,26,32^.

Whilst our data demonstrate that TPM3 expression in hypoxia is controlled by HIF-1 it should be noted that this does not conclusively demonstrate a direct role for HIF-1 or rule out a role for HIF-2 in some contexts. Interestingly, a previous study identified a HIF-1α binding peak 20-30 kb upstream of the TPM3 transcription start site in hypoxic colorectal cancer cell lines (HCT116 and RK0) which overlapped with H3K27ac suggesting a possible distal hypoxia-responsive regulatory element ^33^. Notably, the previously reported proteomic analysis which determined a direct interaction between TPM3 and RIOK3 suggests that in addition to the mechanism of hypoxia-induction described here, TPM3 could also be phosphorylated and potentially stabilised by RIOK3 in hypoxia ^15^.

A previous report has shown that TPM3 mRNA is included in EVs generated by platelet cells therefore raising the question of whether this also occurs in TNBC cells and increases in hypoxia ^14^. Our study is the first to experimentally validate TPM3 as an EV cargo protein. Notably, this is supported by previous proteomic datasets that also detected TPM3 in EV preparations, although these studies were not focused on TPM3 and did not pursue validation or functional investigation ^34,35^. Although we have identified TPM3 as an EV cargo protein we are not able to conclusively conclude that the TPM3 in hypoxic-EVs is responsible for the phenotype observed in normoxic recipient cells. It is also possible that hypoxia-induced TPM3 influences EV content more broadly and the effect observed here is indirect. It is important to note that hypoxic EVs have been described to contain many proteins and micro RNAs which impact invasion, migration, proliferation, angiogenesis and immunomodulation ^36^.

In conclusion, our study identifies TPM3 as a novel, HIF-1–regulated effector that links hypoxia to cytoskeletal remodelling, motility, and intercellular communication in TNBC. By demonstrating its incorporation into EVs and its functional contribution to both hypoxic and normoxic cell behaviour, we highlight TPM3 as a mediator that bridges the heterogeneity of the tumour microenvironment. Together, these findings position TPM3 as both a biomarker of hypoxic adaptation and a promising therapeutic target to constrain TNBC aggressiveness and improve patient outcomes.

## Methods

### Cell lines and reagents

MDA-MB-231 (TNBC, provided by Dr Amanda Coutts, University of Oxford), MDA-MB-453 (TNBC, provided by Dr Isabel Pires, University of Manchester) and BT-549 (TNBC, provided by Prof. Katherine Vallis, University of Oxford). Colorectal RKO and RKO^HIF-1^^α-/-^ cancer cells were grown in DMEM ^37^. Cell culture media was supplemented with 10% FBS and cells were maintained in an incubator set at 37LC and 5% CO₂. All cell lines were verified mycoplasma free using a MycoAlert^TM^ mycoplasma detection kit (Lonza). Inhibitors/drugs used were ATM-3507 (Sigma-Aldrich, 1861449-70-8), Dynasore (Med Chem Express, 304448-55-3). For siRNA-mediated knockdowns, MDA-MB-231 cells were transfected with siRNA to a final concentration of 50 nM using Lipofectamine RNAiMAX (Invitrogen) following the manufacturer’s protocols. siRNA sequences are provided in **Table SI**.

### Hypoxia

A Bactron II anaerobic chamber (Shel Labs) was used for hypoxic treatment at <0.1% O_2_. A Whitley M35 Workstation (Don Whitley Scientific) was used for 2% O_2_ or cyclic hypoxia. Cycling conditions were <0.1% O_2_ for 2 h followed by 2% O_2_ for 2 h as described previously^19^. For all experiments except MTT assays, cells were seeded on glass dishes and harvested inside the chambers with equilibrated reagents.

### Western blotting

Samples were lysed in UTB (9 M Urea and 75 mM Tris-HCl pH 7.5 supplemented with 0.15 M β-mercaptoethanol prior to use). EV samples were lysed in 1× RIPA lysis buffer (Millipore, 20-188) supplemented with protease inhibitor cocktail (Roche, 11873580001). After a brief sonication, proteins were separated on a 4-20% polyacrylamide gel (Bio-Rad) and transferred onto a nitrocellulose membrane (Bio-Rad). Primary antibodies used were: TPM3 (Abcam, ab113692), β-actin (Santa Cruz, sc-69879), HIF-1α (BD Biosciences, 610958), GRP78 (BD Biosciences, 610979), KAP1 (Bethyl, A300-274A), KAP1-S824 (Bethyl, A300-767A), ALIX (Abcam, ab275377), PRDX3 (Abcam, ab73349), GM130 (Abcam, ab52649).

Secondary antibodies: IRDye 680RD goat anti-mouse IgG (LI-COR, 926-68070), IRDye 800CW Goat anti-rabbit IgG (LI-COR, 926-32211), Goat anti-mouse IgG HRP (Invitrogen, 31430), Goat anti-rabbit IgG HRP (Invitrogen, 31460). Images were acquired by chemiluminescence using Odyssey Infrared Imaging (LI-COR Biosciences) or ChemiDoc XRS+ Gel Imaging System (Bio-Rad).

### RT-qPCR

Trizol (Invitrogen) was used to isolate RNA, and the Verso enzyme kit (Thermo Fisher Scientific) to reverse transcribe RNA. SYBR Green PCR Master Mix kit (Applied Biosystems) was used, and the reaction was carried out on a StepOne Real-Time PCR System (Thermo Fisher Scientific) with v2.0.5 software (Applied Biosystems). RNA fold change was measured using a 2^-ΔΔCt^ method relative to the 18S endogenous control gene. Data shown are the mean of three biological replicates ± SEM. All primers sequences are available in supplementary **Table S2**.

### Immunofluorescence

Cells were seeded onto autoclaved cover slips (Menzel-Glaser). Cells were fixed in 4% (w/v) paraformaldehyde in PBS. Samples were permeabilised in 0.1% PBS-Triton X-100 and blocked with 5% (w/v) BSA (Thermo Fisher) in PBS. A LSM710 confocal microscope (Carl Zeiss Microscopy Ltd) was used for imaging. Antibodies/reagents used: TPM3 (Abcam, ab113692), β-actin (Santa Cruz, sc-69879), Alexa Fluor 488-conjugated goat anti-mouse (Invitrogen, A32723), Alexa Fluor Plus 647 Phalloidin (Invitrogen, A30107).

### 3-(4,5-dimethylthiazol-2-yl)-2,5-diphenyl-2H-tetrazolium bromide (MTT) assay

Cells were seeded in flat bottomed 96-well plates. Each experiment was carried out in triplicate. MTT reagent (5 mg/mL, Invitrogen) was added to cells in the dark for 3 hours (37°C, 5% CO_2_). After MTT containing culture media was removed, DMSO was added to cells and incubated in the dark for 15 minutes (37°C). The plate was then read using a POLARstar Omega plate reader (BMG Labtech) (absorbance 570 nm). Cell viability was measured relative to the untreated control.

### Flow cytometry

Cells were fixed in ice-cold 70% ethanol. Propidium Iodide (Sigma Aldrich) and RNaseA (NEB) were added to samples for 15 minutes incubation (37°C). Samples were run on a CytoFLEX flow cytometer (Beckman Coulter Life Science) and results were analysed with FloJo software.

### Wound healing assay

Cells were seeded and grown to 95-100% confluency before treatment. At least five parallel wounds were made by scratching the cell monolayer with a 20 μL pipette tip. After rinsing in PBS, the cells were incubated in 0.5% FBS containing culture media. EVOS M5000 (Thermo Fisher) was used to image the wounds immediately after scratching and over time. The area of each wound was measured by ImageJ (National Institutes of Health). The wound closure % was calculated using the following formula where *A* is the area of the wound:

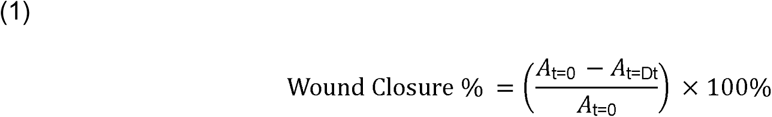

### Invasion assay

Cells were seeded in an 8 μm pore size BioCoat^TM^ Matrigel Invasion Chamber (Corning). Trypsinised cells (5×10^5^) were added to the upper chambers of a 24-well plate in DMEM containing 0.1% FBS. After treatment, the motile cells at the bottom of the filter were stained with crystal violet (0.5% w/v in 50% MeOH and 20% EtOH). The number of cells which had invaded was measured by counting the stained cells using EVOS M5000 (Thermo Fisher).

### Colony survival assay

Cells were seeded at the appropriate density for siRNA transfection/drug treatment under normoxia or hypoxia for the indicated periods, each experiment was carried out in triplicate. Colonies were allowed to grow for 8-10 days in a standard humidified incubator at 37°C and 5% CO_2_. Once the colonies formed (≥50 cells), crystal violet (0.5% w/v in 50% MeOH and 20% EtOH) was used for staining. Colonies were counted using an automated colony counter GelCount^TM^ (Oxford Optronix) with GelCount (version 1.2) software, or a manual cell counter (Stuart Scientific). The survival fraction was calculated by number of colonies counted/number of cells seeded × PE, where PE is the plating efficiency of the untreated control (number of colonies counted/number of cells seeded). Colonies with radiation treatment in hypoxic conditions were carried out as previously described ^38^.

### Transmission electron microscopy (TEM)

Cells were seeded onto glass coverslips and cultured until approximately 70% confluency was reached. Following treatment, cells were fixed in 2.5% glutaraldehyde and 4% PFA in

0.1 M PIPES buffer (pH 7.2) followed by washes in 0.1 M PIPES buffer, incubated in 50 mM glycine/PIPES for 15 minutes, washed once in 0.1 M PIPES, embedded in low-melting-point agarose, chilled, trimmed into 1–2 mm blocks, and returned to buffer. Samples were then treated with 1% osmium tetroxide and 1.5% potassium ferrocyanide in 0.1 M PIPES buffer for 1 hour at 4°C. Samples were washed in Milli-Q water and incubated overnight in 0.5% uranyl acetate at 4°C in the dark, followed by washes in Milli-Q water. Dehydration was performed on ice using a graded ethanol series (30%, 50%, 70%, 80%, 90%, 95%, and 3 × 100%) at room temperature. Samples were infiltrated with Taab low-viscosity epoxy resin via ethanol:resin series (3:1 for 1 hour, 1:1 for 1.5 hours, and 1:3 for 1 hour), then 100% resin at room temperature overnight. Resin was refreshed twice the next day, with centrifugation (12,000 rpm, 3 minutes) between changes. Agarose-embedded blocks were transferred to Beem capsules containing fresh resin and polymerised at 60°C for a minimum of 24 hours. Ultrathin sections (∼90 nm) were prepared using a Diatome diamond knife on a Leica UC7 ultramicrotome, mounted onto 200 mesh copper grids, and post-stained using Reynolds’ lead citrate for 5 minutes at room temperature, followed by washes in Milli-Q water. Sections were imaged using a JEOL 1400 transmission electron microscope equipped with a Gatan Rio CMOS detector.

### EV isolation and purification

After treatment, culture media (FBS-free DMEM) was collected and transferred to a 50 mL Falcon tube for EV isolation. EV-containing media was centrifuged at 111.8 × g for 5 minutes at 4°C (Jouan CR4i Centrifuge, Thermo Electron Corporation). Supernatant was collected, transferred to a new Falcon tube, and centrifuged at 1,006.2 × g for 10 minutes at 4 °C. Supernatant was collected, transferred to a new Falcon tube and centrifuged at 1,788.8 × g for 30 minutes at 4°C. The supernatant was transferred to an ultracentrifuge tube (Ultra-Clear centrifuge tubes, Beckman Coulter). Ultracentrifugation was conducted in a pre-cooled SW 32.1 Ti Swinging-Bucket rotor in an Optima XPN-80 Ultracentrifuge (Beckman Coulter) at 87,945 × g for 140 minutes at 4°C. Following ultracentrifugation, the media was removed, and the pellet resuspended in cold, sterile PBS. Samples were ultracentrifuged again at 87,945 × g for 140 min at 4°C. Supernatant was carefully aspirated leaving the purified EVs, which were resuspended in sterile PBS or fresh 10% FBS supplemented DMEM for subsequent experiments.

### Nanoparticle Tracking Analysis (NTA)

Isolated EVs were resuspended in equal volumes of 1 × PBS across all conditions. EV number and size distribution were assessed using ZetaView® (Particlemetrix) according to the manufacturer’s instructions. Stock EV samples were diluted in 1 × PBS to working concentrations ranging from 1:10,000 to 1:100,000 and loaded into the flow cell. Measurements were recorded as videos from 8-11 separate positions across the flow cell. EV number and size were estimated from this using the ZetaView analysis software

### Statistical analysis

A two-tailed, paired Student’s *t*-test was used for the comparison of two means and a two-way analysis of variance (ANOVA) with Tukey’s multiple comparisons or Šídák’s multiple comparison test were used for the comparison of more than two means.

## Acknowledgements

We are grateful to Raman Dhaliwal and Dr Charlotte Melia at the Sir William Dunn School of Pathology Electron Microscopy Facility. CZ was supported by Breast Cancer Now (2022FebPR1492 awarded to EMH and EEP). SAT and EMH thank the EPSRC for the support of programme grant EP/S019901/1. GB was supported by a UNIQ+ Research Internship (University of Oxford). EP was supported by a Clarendon scholarship (University of Oxford).

## Contributions

CZ, JTC, KF, PS, ST, EP, GB, EEP and EMH were responsible for collecting, analysing, and interpreting data. EMH and CZ drafted the manuscript. EMH was responsible for study conceptualisation and study oversight. JTC, CZ and EMH were responsible for figure creation. All authors critically reviewed the manuscript and approved the final version.

## Data Availability

All data supporting the findings of this study are available within the paper and its Supplementary Information or by request to the corresponding author.

