## Supplementary material for "HIF-1–regulated TPM3 links hypoxia to motility and invasion beyond the hypoxic fraction in triple-negative breast cancer": SI

Contents | Page number

| Contents | Page number |
|---|---|
| SI Tables | 2 |
| SI Figures | 3 |
| Uncropped blots for main figures | 11 |

SI Tables

**Table S1.** Sequences of siRNA

| Gene | Sequence | Working concentration | Source |
|---|---|---|---|
| TPM3 | AGGUAUGAAGGUUAUUGAA [dT][dT] | 50 nM | Sigma Aldrich |
| TPM3 | GAGAGAGGUAUGAAGGUUA | 50 nM | Dharmacon |
| TPM3 | GAAGAGGCAGAUAGGAAGU | 50 nM | Dharmacon |
| TPM3 | GAUUCUUACUGAUAAACUC | 50 nM | Dharmacon |
| TPM3 | UGAGUUUGCUGAGAGAUCG | 50 nM | Dharmacon |

**Table S2**. Sequences of primers used for RT-qPCR.

| Gene | Oligonucleotide name | Sequence | Working concentration | Source |
|---|---|---|---|---|
| 18S | 18S_F<br>18S_R | TAGAGGGACAAGTGGCGTTC<br>CGGACATCTAAGGGCATCAC | 10 μM | Sigma Aldrich |
| TPM3 | TPM3_F1<br>TPM3_R1 | ACCACCATCGAGGCGGTAA<br>CCCTTTCCTCCGCATCATCA | 5 μM | Sigma Aldrich |
| TPM3 | TPM3_F2<br>TPM3_R2 | ATCCAGCTGGTTGAAGAGGA<br>CACCAACTTACGAGCCACCT | 5 μM | Sigma Aldrich |
| TPM3 | TPM3_F3<br>TPM3_R3 | CTCAAGTCTCTTGAGGCTCAGG<br>TTCCAGCTTGGCTACCGATCTC | 5 μM | Sigma Aldrich |

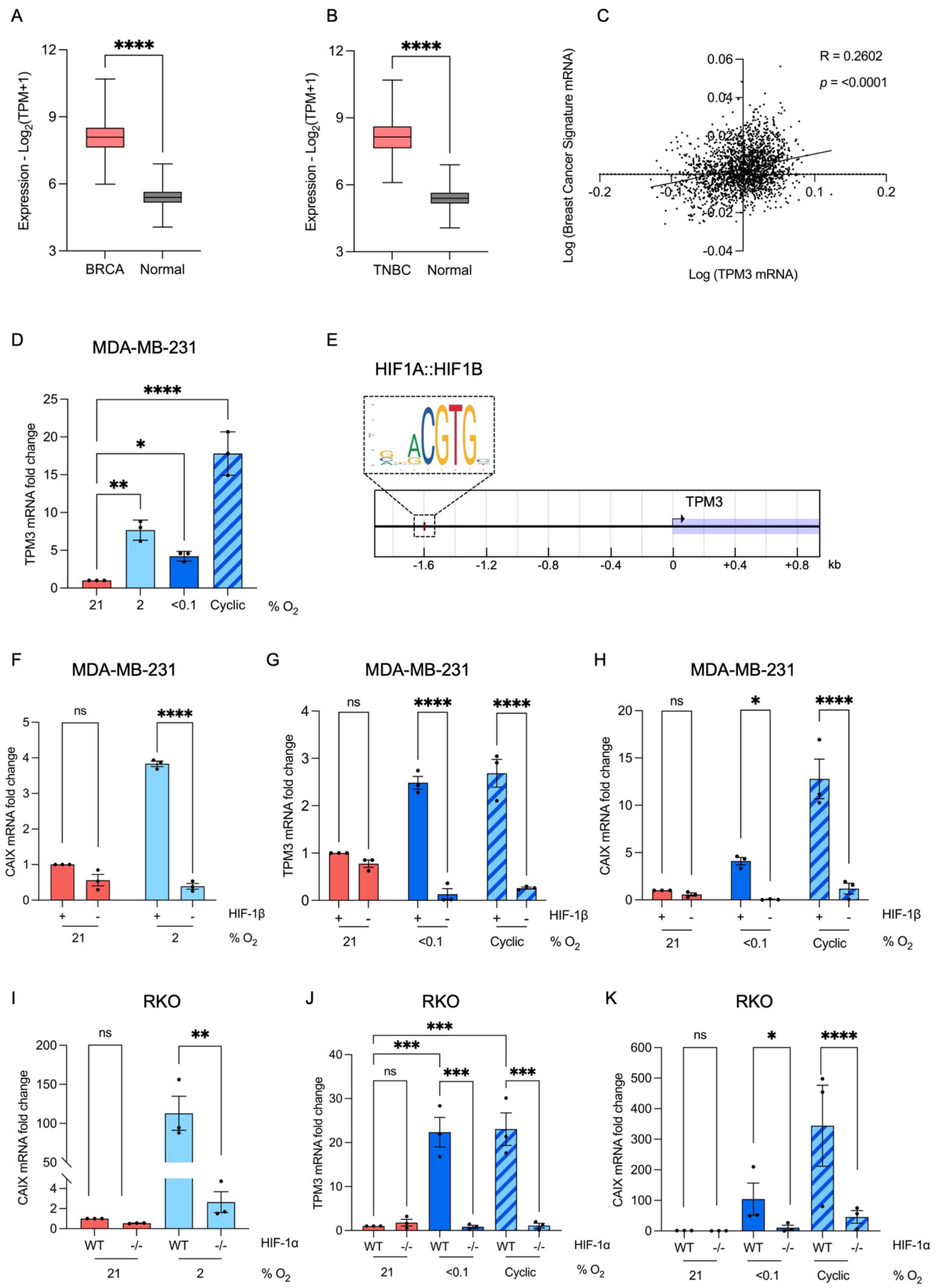


**Figure S1. TPM3 is induced in a HIF-1 dependent manner in a range of hypoxic conditions A.** TPM3 mRNA levels in BRCA and normal breast tissue generated using TPM3 mRNA expression from Metabric dataset. num(BRCA)=1980; num(Normal)=514. **B.** TPM3 mRNA levels in TNBC and normal breast tissue generated using TPM3 mRNA expression from Metabric. num(TNBC)=335; num(Normal)=514. **C.** Correlation between the BRCA-specific hypoxia signature [1] and TPM3. TPM3 mRNA expression from Metabric dataset.

Statistical analysis was determined using simple linear regression test and Pearson correlation. **D.** MDA-MB-453 cells were exposed to a range of hypoxic conditions for 16 h followed by RT-qPCR for TPM3. Statistical testing was done using a paired *t*-test. **E.** The positions of a hypoxia-responsive element (HRE) was determined using the Eukaryotic Promoter Database (p-value cutoff, 0.0001). The illustrated transcription factor binding motifs are from the JASPAR CORE database [2,3]. **F.** MDA-MB-231 cells were treated with siHIF-1β or scramble control then exposed to 21% or 2% $O_2$ for 16 h, followed by RT-qPCR for CAIX. Statistical testing was done using a paired *t*-test. **G.** MDA-MB-231 cells were treated with siHIF-1β or scramble control followed by exposure to the indicated oxygen level for 16 h, TPM3 mRNA level was determined and shown relative to the normoxic control. Normoxic data is the same as shown in Figure 1H. Statistical testing was done using a paired *t*-test. **H.** MDA-MB-231 cells were treated with siHIF-1β, or scramble control followed by exposure to the indicated oxygen level for 16 h, CAIX mRNA level was determined and shown relative to the normoxic control. Normoxic data is the same as shown in part E. Statistical testing was done using a paired *t*-test. **I.** RKO and $RKO^{HIF\text{-}1\alpha^{-/-}}$ cells were exposed to 21% and 2% $O_2$ for 16 h, followed by RT-qPCR for CAIX. Statistical testing was done using a paired *t*-test. **J.** RKO and $RKO^{HIF\text{-}1\alpha^{-/-}}$ cells were exposed to the indicated hypoxic conditions for 16 h followed by RT-qPCR for TPM3. Normoxic data is the same as shown in Figure 1I. Statistical testing was done by two-way ANOVA with Tukey's multiple comparisons test. **K.** RKO and $RKO^{HIF\text{-}1\alpha^{-/-}}$ cells were exposed to the indicated oxygen level for 16 h, followed by RT-qPCR for CAIX mRNA. Normoxic data is the same as shown in Figure 1I. Statistical testing was done using a paired *t*-test. Data shown from three separate experiments ($n$ = 3) are displayed ± standard error of the mean (SEM). * $p < 0.05$, ** $p < 0.01$, *** $p < 0.001$ and ns $p > 0.05$.

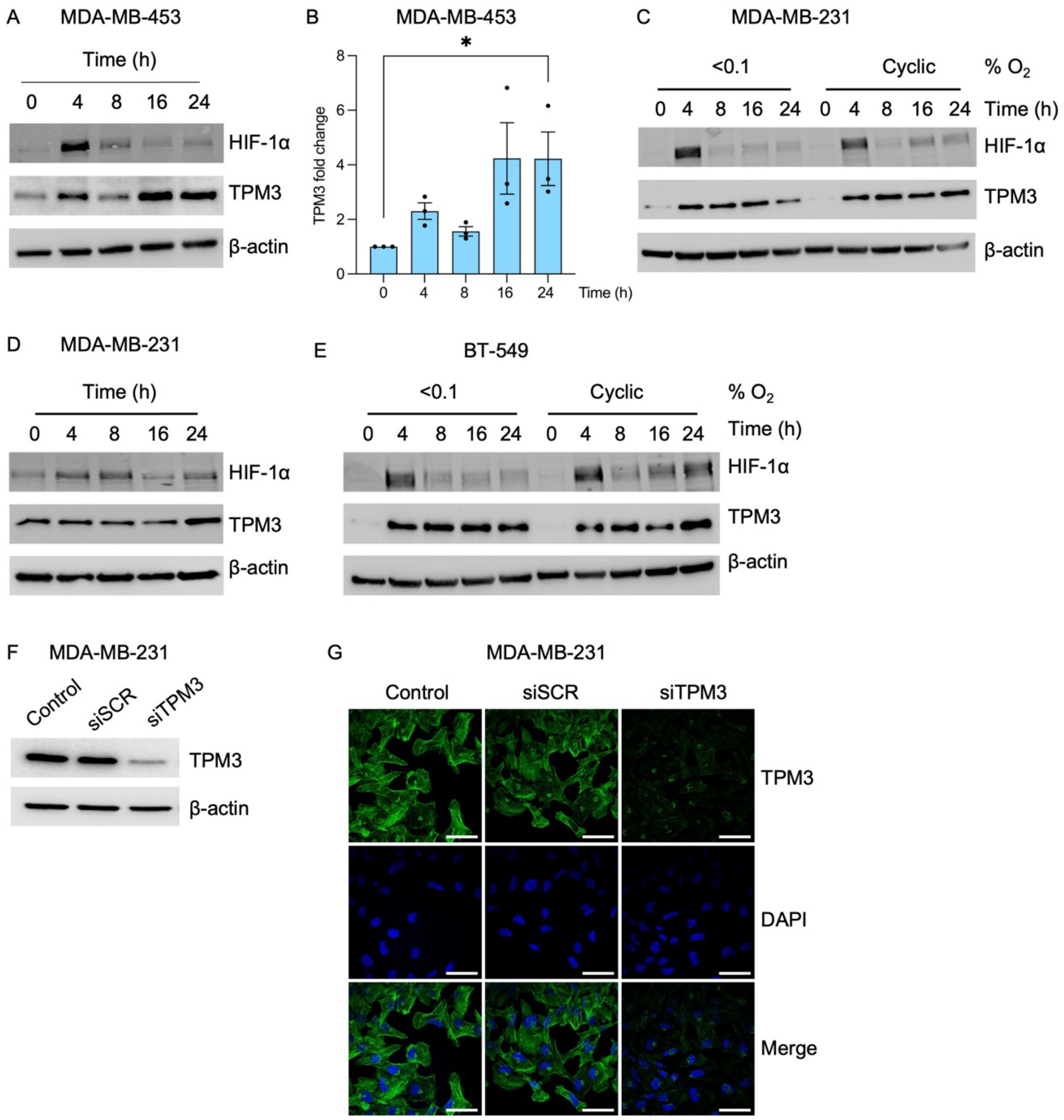


**Figure S2. TPM3 is induced at the protein level in hypoxia A.** MDA-MB-453 cells were exposed to 2% $O_2$ for the indicated times, followed by western blotting for proteins shown. **B**. Quantification of result A. Statistical testing was done by paired *t*-test. **C.** MDA-MB-231 cells were exposed to <0.1% $O_2$ and cyclic hypoxia for the indicated times, followed by western blotting for proteins shown. **D.** BT-549 cells were exposed to 2% $O_2$ for the indicated times, followed by western blotting for proteins shown. **E.** BT-549 cells were exposed to <0.1% $O_2$ and cyclic hypoxia for the indicated times, followed by western blotting for proteins shown. **F.** MDA-MB-231 cells were treated with siRNA to TPM3, or scramble control (siSCR) followed by western blotting for protein indicated. **G.** Representative immunofluorescence images confirming TPM3 knockdown. TPM3 (green), DAPI (blue). Scale bar is 50 μm. Cells collected from the same experiment in part F. Data shown from three separate experiments ($n$ = 3) are displayed as mean ± SEM. * $p < 0.05$, ** $p < 0.01$, *** $p < 0.001$ and ns $p > 0.05$.

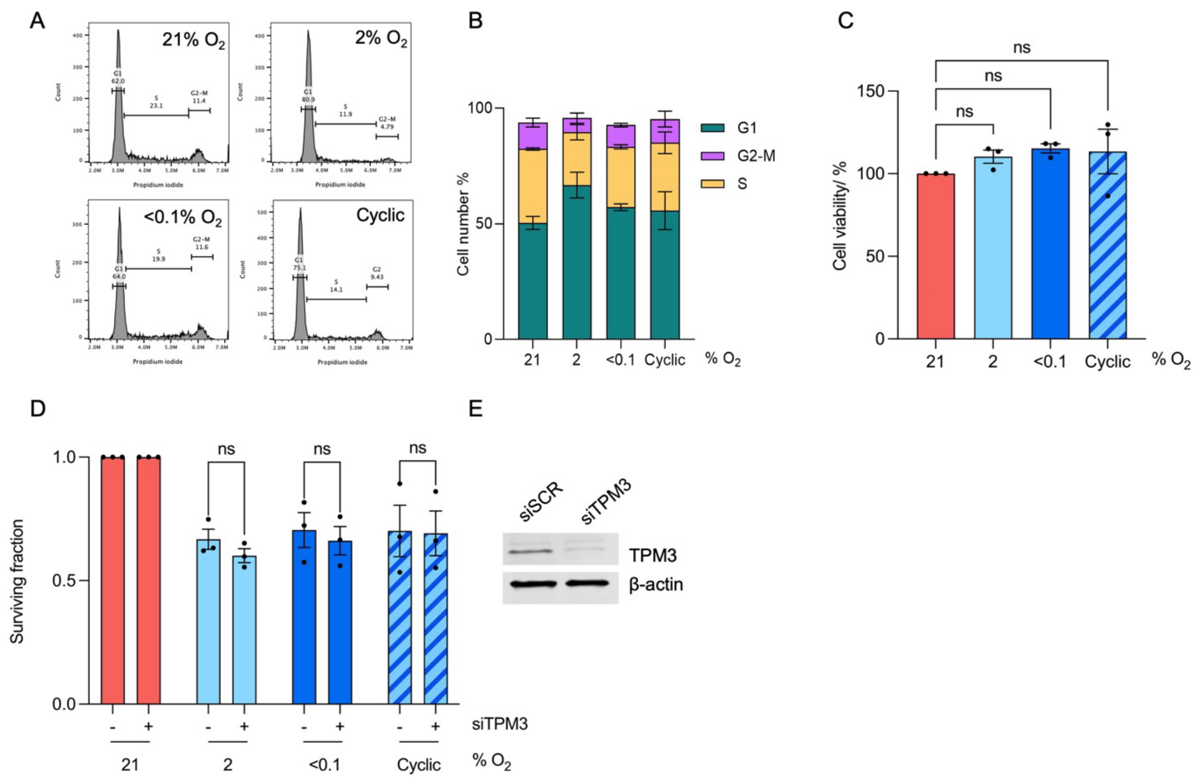


**Figure S3. The range of hypoxic conditions used or loss of TPM3 did not cell viability** **A.** MDA-MB-231 cells were exposed to the indicated oxygen concentrations for 16 h, propidium iodide (PI) was added to sample to determine total DNA content prior to collection and analysed by flow cytometry. **B.** Quantification of part A, indicating the percentage of cell number in each condition. No statistically significant differences were observed, as determined by two-way ANOVA, Tukey's multiple comparisons test. **C.** MDA-MB-231 cells were exposed to the indicated hypoxia treatments for 16 h, cell viability was measured using the MTT assay. Statistical testing was done by paired *t*-test. **D.** MDA-MB-231 cells were treated with siTPM3 or a scramble siRNA and then exposed to the indicated oxygen concentrations for 16 h, followed by a colony survival assay. Statistical testing was done by paired *t*-test. **E.** MDA-MB-231 cells were treated with siTPM3 or a scramble siRNA (siSCR) followed by western blotting for TPM3 and β-actin as a loading control. Data shown from three separate experiments ($n$ = 3) are displayed mean ± SEM. * $p < 0.05$, ** $p < 0.01$, *** $p < 0.001$ and ns $p > 0.05$.

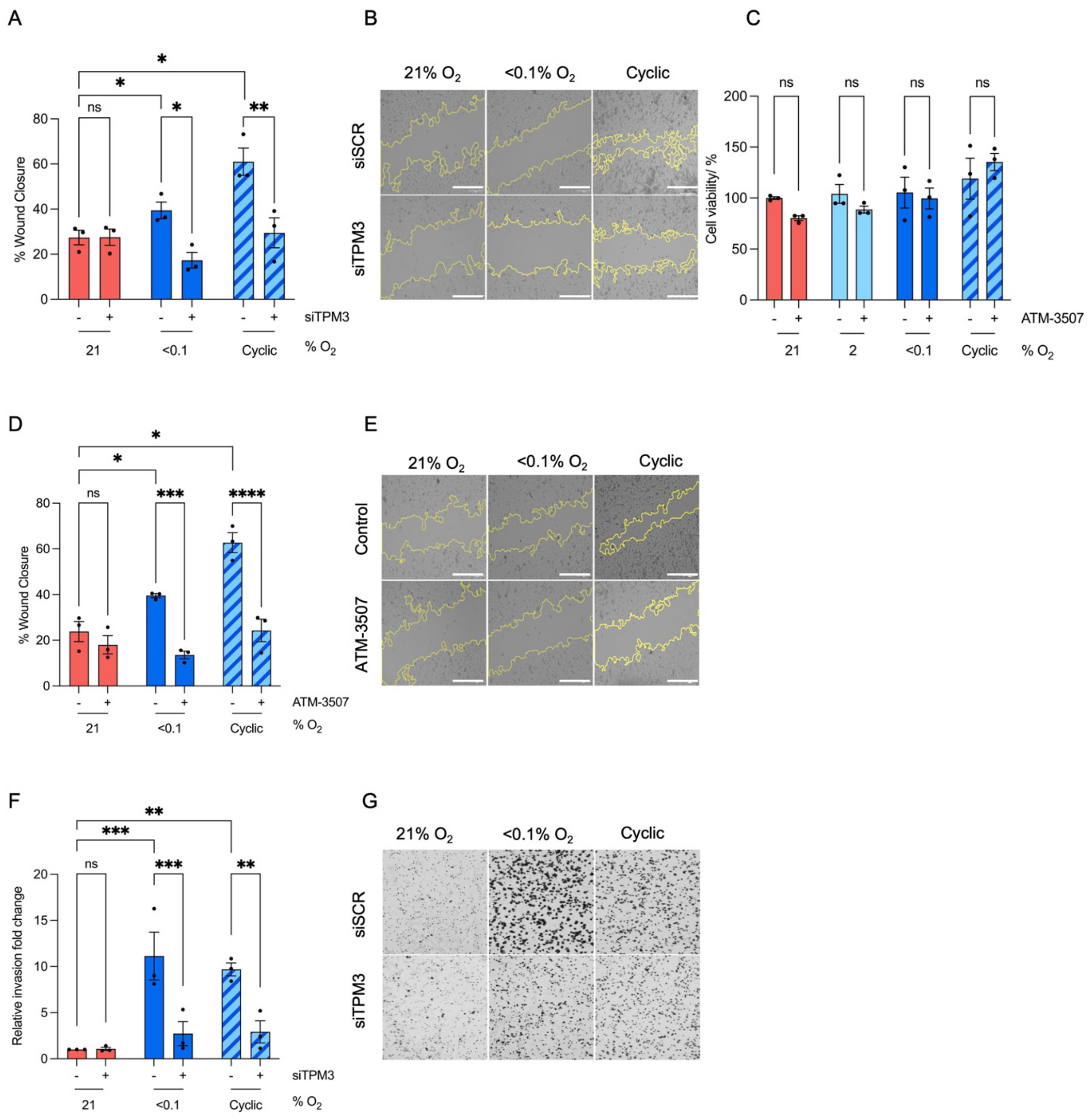


**Figure S4. Hypoxia-mediated cell motility is TPM3-dependent A.** MDA-MB-231 cells were treated with siTPM3 or a scramble siRNA followed by exposure to the indicated oxygen conditions for 8 h. The % of wound closure is shown in each condition. Normoxic data are the same as shown in Figure 3A. Statistical testing was done by two-way ANOVA with Šídák's multiple comparison test. **B.** Representative images of wound healing assay in part A. Scale bars are 200 µm. **C.** MDA-MB-231 cells were treated with ATM-3507 (6 µM) for 1 h prior to and during exposure to the indicated oxygen conditions for 16 h, cell viability was measured using the MTT assay. Statistical testing was done by paired *t*-test. **D.** MDA-MB-231 cells were treated with ATM-3507 (6 µM) for 1 h prior to and during exposure to indicated hypoxic conditions for 8 h. The % of wound closure is shown in each condition. Normoxic data are the same as shown in Figure 3D. Statistical testing was done using a two-way ANOVA with Šídák's multiple comparison test. **E.** Representative images of wound healing assay in part D. Scale bars are 200 µm. **F.** MDA-MB-231 cells were treated with siTPM3 or a scramble siRNA followed by exposure to the indicated oxygen concentrations for 16 h. The invasion fold change relative to normoxic control is shown in each condition. Normoxic data are the same as shown in Figure 3E. Statistical testing was done by two-way ANOVA with Šídák's multiple comparison test. **G.** Representative images of wound healing assay in part F. Data from three separate experiments ($n = 3$) are displayed as mean ± standard error of the mean (SEM) unless specified otherwise. * $p < 0.05$, ** $p < 0.01$, *** $p < 0.001$ and **** $p < 0.0001$.

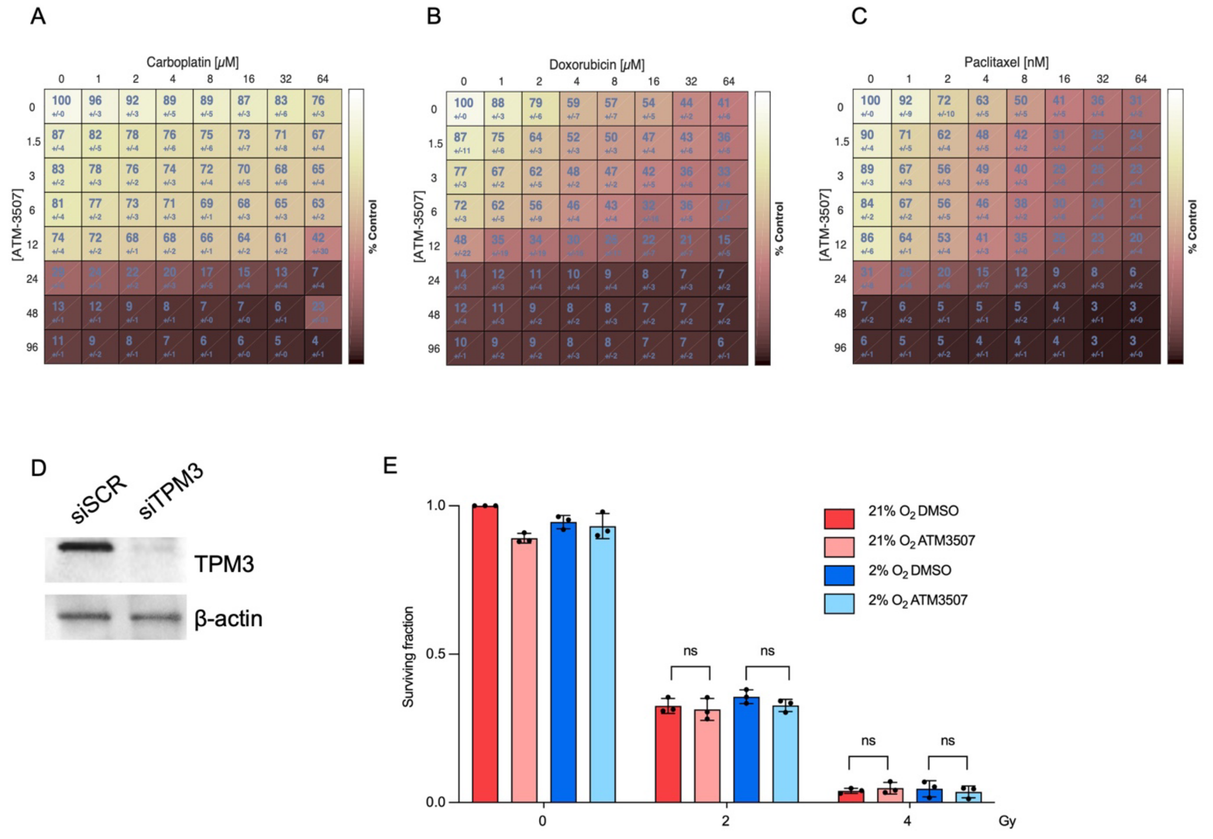


**Figure S5. Inhibition or loss of TPM3 combines well with standard of care for TNBC**
**A**. MDA-MB-231 cells were treated with Carboplatin (1 - 64 μM) for 48 h and ATM-3507 (1.5-12 μM) for 17 h. For the 16 h before an MTT assay was carried out the cells were in hypoxia (2% $O_2$). **B**. MDA-MB-231 cells were treated with Doxorubicin (1 - 64 μM) for 24 h and ATM-3507 (1.5-12 μM) for 17 h. For the 16 h before an MTT assay was carried out the cells were in hypoxia (2% $O_2$). **C**. MDA-MB-231 cells were treated with Paclitaxel (1 - 64 nM) for 72 h and ATM-3507 (1.5-12 μM) for 17 h. For the 16 h before an MTT assay was carried out the cells were in hypoxia (2% $O_2$). In A, B and C cell viability score was assessed by an interactive platform Combenefit. Dose-response matrices with drug concentrations on axes and combination effects as heatmap overlay are shown. White shading indicates higher cell viability and brown shading indicates reduced cell viability. Statistical testing was done using the built-in analysis algorithm [4]. Data from three separate experiments ($n$ = 3). * $p < 0.05$, ** $p < 0.01$, *** $p < 0.001$ and **** $p < 0.0001$. **D**. Validation by western blotting that siTPM3 in Figure 4D was effective. **E**. MDA-MB-231 cells were treated with ATM-3507 (6 μM) for 1 h prior to and during exposure to 21% or 2% $O_2$ for 16 h prior to irradiation (0, 2, 4 Gy). Cells in hypoxia were irradiated in hypoxic conditions. After irradiation, all cells were returned to 21% $O_2$ and a colony survival assay carried out. Statistical testing was done by paired t-test. Data from three separate experiments (n = 3) are displayed with mean ± standard error of the mean (SEM).

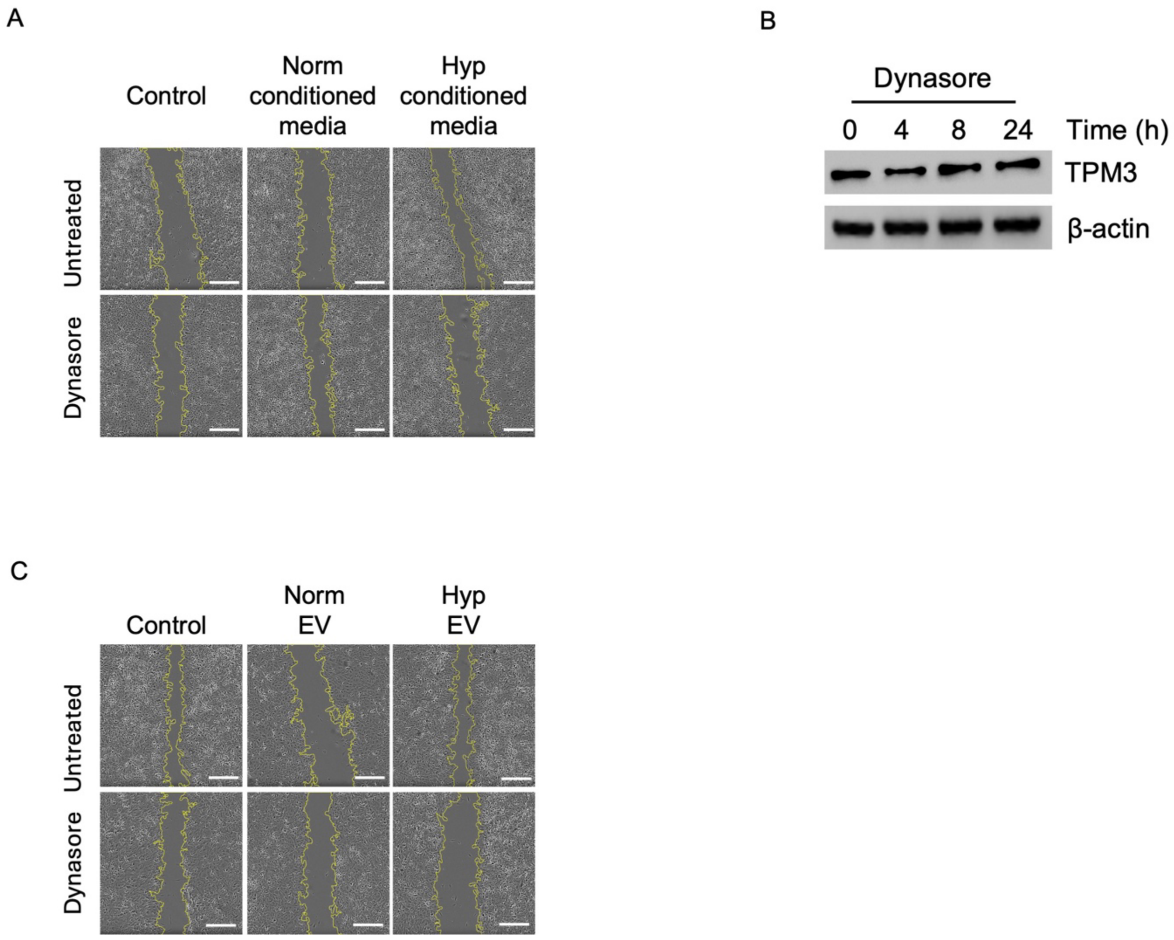


**Figure S6. EVs generated in hypoxic cells drive motility of normoxic cells in a TPM3-dependent manner A.** Representative images of wound healing assay from Figure 5G. Scale bars are 500 µm. **B.** MDA-MB-231 cells were treated with Dynasore (50 µM) and harvested at the indicated times followed by western blotting for the proteins shown. **C.** Representative images of wound healing assay from Figure 6G. Scale bars are 500 µm.

### Uncropped blots

#### Figure 1E

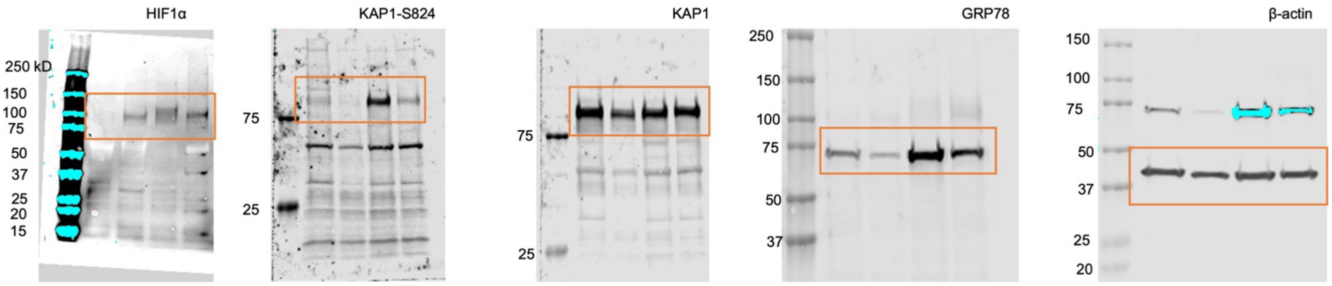


#### Figure 1J

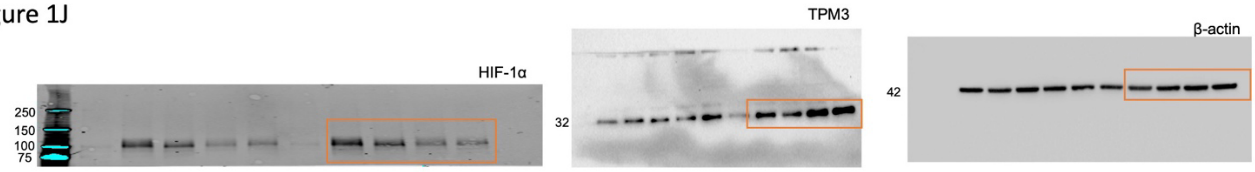


#### Figure 3B

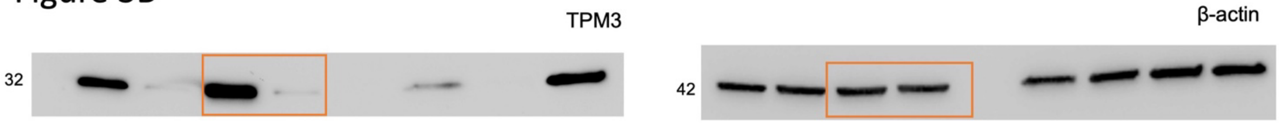


#### Figure 3G

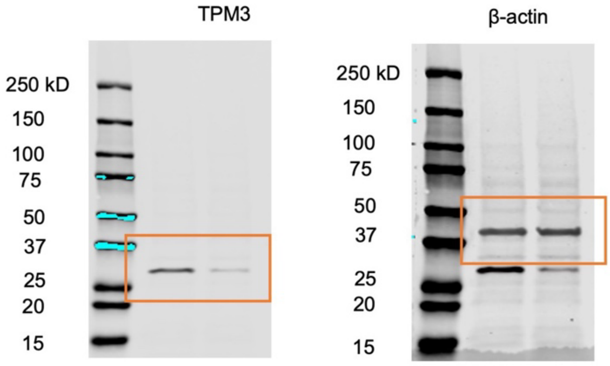


#### Figure 5B

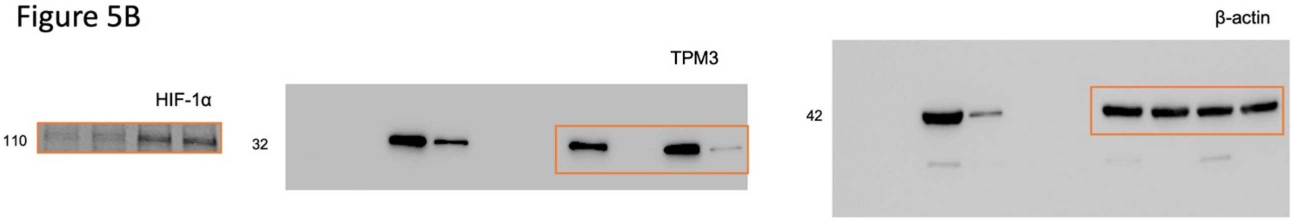

Figure 5E

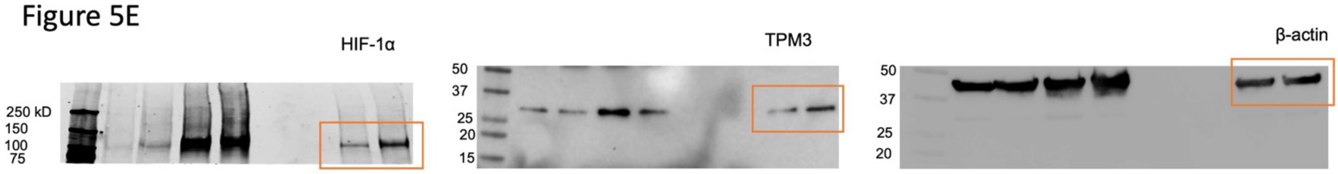


### Figure 6E

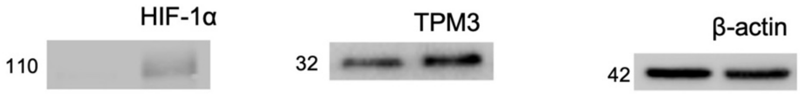


### Figure 7C

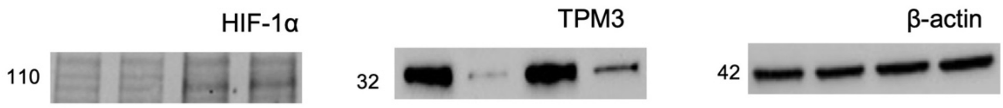


Figure 7D

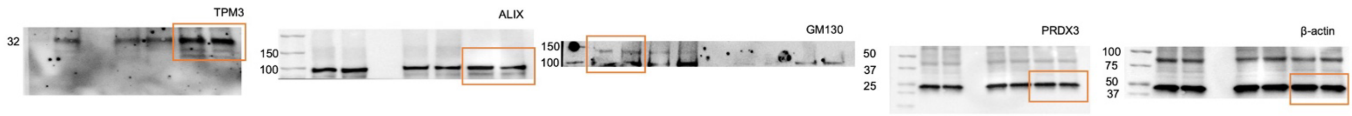


Figure 7E

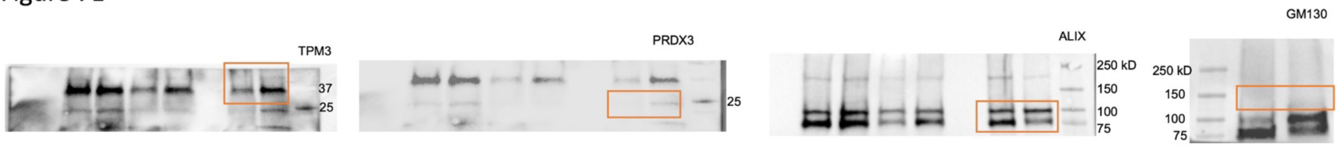
